# Structural modeling reveals viral proteins that manipulate host immune signaling

**DOI:** 10.1101/2025.07.12.664507

**Authors:** Nitzan Tal, Romi Hadari, Renee B. Chang, Ilya Osterman, Roy Jacobson, Erez Yirmiya, Nathalie Bechon, Dina Hochhauser, Miguel López Rivera, Barak Madhala, Jeremy Garb, Tanita Wein, Philip Kranzusch, Gil Amitai, Rotem Sorek

## Abstract

Immune pathways that use intracellular nucleotide signaling are common in animals, plants and bacteria. Viruses can inhibit nucleotide immune signaling by producing proteins that sequester or cleave the immune signals. Here we analyzed evolutionarily unrelated signal-sequestering viral proteins, finding that they share structural and biophysical traits in their genetic organization, ternary structures and binding pocket properties. Based on these traits we developed a structure-guided computational pipeline that can sift through large phage genome databases to unbiasedly predict phage proteins that manipulate bacterial immune signaling. Numerous previously uncharacterized proteins, grouped into three families, were verified to inhibit the bacterial Thoeris and CBASS signaling systems. Proteins of the Sequestin and Lockin families bind and sequester the TIR-produced signaling molecules 3′cADPR and His-ADPR, while proteins of the Acb5 family cleave and inactivate 3′3′-cGAMP and related molecules. X-ray crystallography and structural modeling, combined with mutational analyses, explain the structural basis for sequestration or cleavage of the immune signals. Thousands of these signal-manipulating proteins were detected in phage protein databases, with some instances present in well-studied model phages such as T2, T4 and T6. Our study explains how phages commonly evade bacterial immune signaling, and offers a structure-guided analytical approach for discovery of viral immune-manipulating proteins in any database of choice.

## Introduction

Intracellular signaling plays a fundamental role in innate immunity across all domains of life. A diverse set of immune pathways, widespread in animals, plants and bacteria, use modified nucleotides as immune signals. These pathways typically include a protein that produces the immune signaling molecule once pathogen invasion is detected, and another protein (or protein complex) that receives the immune signal and triggers a downstream immune response^1–3^. A prototypic such pathway is the human cGAS-STING pathway, in which the cGAS protein produces 2′3′ cyclic GMP-AMP (cGAMP) in response to cytoplasmic DNA, and the STING protein activates immunity once triggered by cGAMP^4^. The cGAS-STING pathway evolved from a bacterial anti-phage defense system called CBASS^5,6^. A wide diversity of CBASS systems was documented in bacteria, each generating a distinct nucleotide immune signal, including cGAMP, cyclic UMP-AMP (cUA), cyclic UMP-UMP (cUU), cyclic AMP-AMP-GMP (cAAG) and many others^5–9^. Another family of bacterial antiviral defense system, Pycsar, produces the molecules cUMP and cCMP for anti-phage immune signaling^10^.

TIR domains were also documented as components of immune signaling systems, particularly in plants and bacteria. Once a protein with a TIR domain senses infection, it produces an immune signaling molecule that contains derivatives of ADP-ribose (ADPR)^1–3^. In bacteria, TIR signaling is employed as part of the Thoeris anti phage defense system. Type I Thoeris systems produce the molecule 3′cADPR^11^ (also called 1′′–3′ gcADPR); type II Thoeris produces a molecule comprising a histidine conjugated to ADPR (His-ADPR)^12^; while type IV Thoeris systems produce the molecule N7-cADPR^13^. These molecules act as second messengers, binding a second protein in the Thoeris pathway to trigger regulated cell death or growth arrest in response to infection. Bacterial Thoeris systems were proposed to be the ancestors of plant TIR-dependent immune pathways that similarly use ADPR derivatives for immune signaling^14^.

As immune signaling pathways are present in a large fraction of bacteria, archaea, plants and animal species^1,8^, viruses evolved anti-defense proteins to eliminate immune signaling molecules from the cell during infection^15^. Some of these anti-defense proteins are enzymes that cleave and inactivate the molecule, including phage-encoded Anti-CBASS1 (Acb1) enzymes that cleave cyclic oligonucleotides produced by CBASS systems^16^, and Anti-pycsar1 (Apyc1) that cleaves cCMP and cUMP to inhibit Pycsar immunity^16^. Pox viruses also encode cGAMP-cleaving enzymes called poxins, allowing virus escape from cGAS-STING immunity^17^.

A second class of viral proteins that inhibit host immune signaling are called “sponge” proteins. These proteins tightly bind and sequester the immune signaling molecules, preventing them from activating downstream immunity^11,18–20^. Sponge proteins were previously discovered in phages, including the Thoeris anti-defense 1 (Tad1) and Tad2 sponges that inhibit bacterial Thoeris systems by binding TIR-produced signaling molecules^11,12,18^, and Acb2 and Acb4 that bind cyclic oligonucleotides to inhibit CBASS ^19,20^.

To date, the identification of viral proteins that target immune signaling molecules has largely depended on serendipity or on experiments with a limited set of viruses^11,18–20^. Here, we identify structural and biophysical features common to viral nucleotide-binding anti-defense proteins, allowing structure-guided discovery of such proteins in large databases of phage genomes. We describe three new families of anti-defense proteins targeting host immune signaling, cumulatively represented in over 5000 proteins within phage protein datasets. We further characterize the interactions of these proteins with their target molecules using in-depth biochemical and structural analyses, providing a molecular explanation for their ability to efficiently overcome bacterial defenses.

## Results

### Protein fusion analysis reveals a new sponge family

While analyzing known viral proteins that inhibit bacterial immune signaling, we observed that proteins from two well-characterized sponge families, Tad2 and Acb4, are frequently encoded as a single fused protein in phage genomes (Fig. 1A). We hypothesized that fusion of two distinct sponge domains can grant phages the ability to simultaneously block multiple immune signaling molecules using a single protein. Indeed, co-expression of an Acb4-Tad2 fusion protein from *Pasteurella phage* Pm86 with either type I Thoeris or with type I CBASS canceled the ability of both these systems to provide defense against phages (Fig S1A,B).

**Figure 1.**
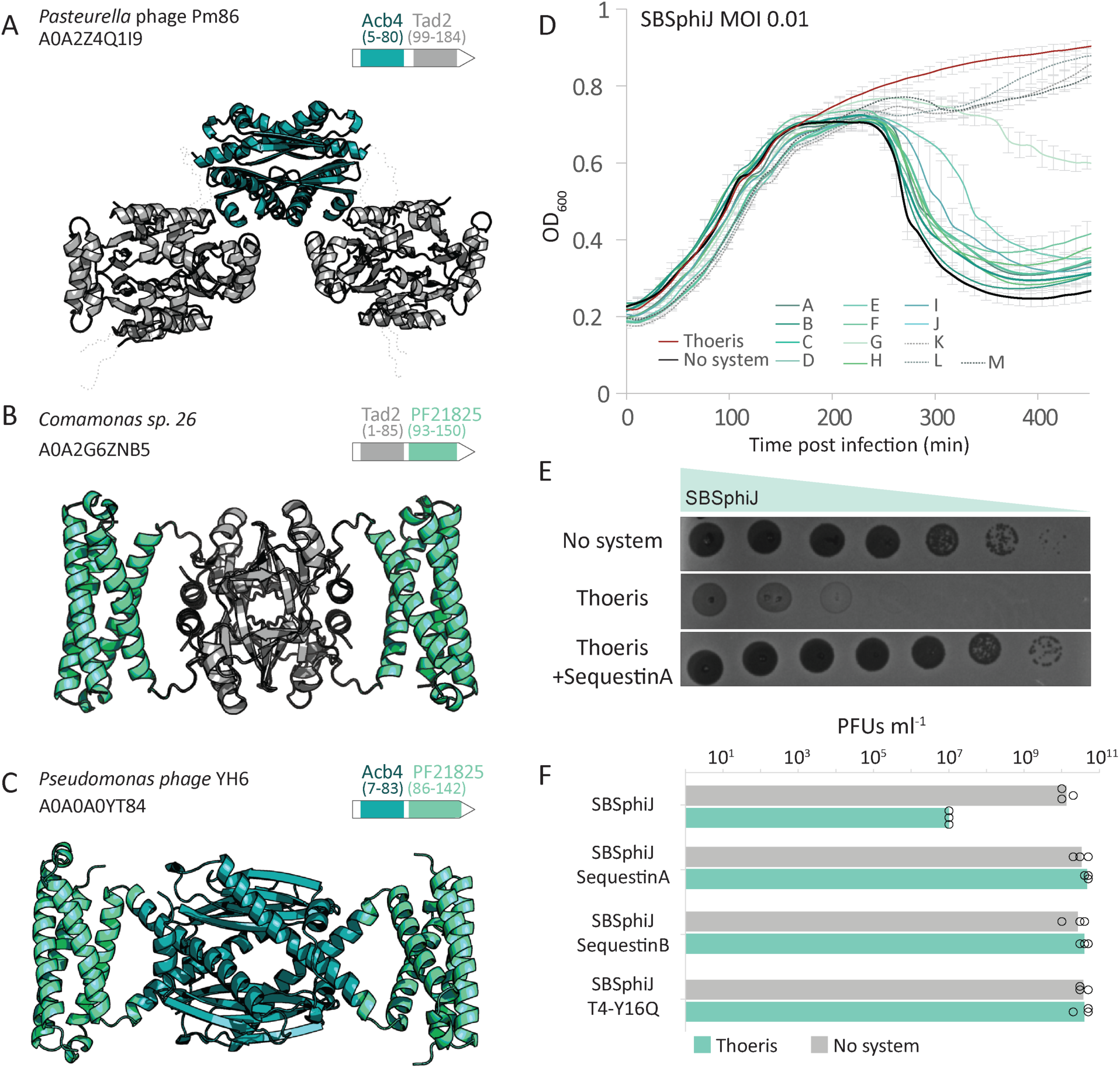
Proteins with PF21825 domain inhibit type I Thoeris defense. (A) A fusion protein of Tad2 and Acb4, found in *Pasteurella* phage Pm86. Each of the Tad2 and Acb4 domains was modeled separately by AlphaFold3 as a homotetramer (Acb4, ipTM = 0.91; Tad2, iPTM = 0.90), and the two models were overlayed. Dashes represent unmodeled linkers between Acb4 and Tad2 domains. (B) A fusion protein of Tad2 and a PF21825 domain, detected in a mobile genetic element in *Comamonas sp*. 26. Model of a homotetramer, ipTM=0.74. (C) A fusion protein of PF21825 domain and Acb4 from *Pseudomonas* phage YH6. Model of a homotetramer, ipTM=0.74. (D) Growth curves of *B. subtilis* cells expressing the type I Thoeris system alone (red), or co-expressing the Thoeris system and a gene with PF21825 pfam domain (A-M represent the 13 tested PF21825 genes, shades of turquoise), or a control strain without a defense system (black), infected by phage SBSphiJ at an MOI of 0.01. Each curve is the average of three replicates, with error bars indicating standard deviation. (E) A representative PF21825 gene (SequestinA, name explained below) that cancels Thoeris defense. Shown are tenfold serial dilution plaque assays, comparing the plating efficiency of phage SBSphiJ on bacteria that express the type I Thoeris defense system alone, or co-expressing the Thoeris system with SequestinA. The control strain lacks the system and expresses GFP instead. Images are representative of three replicates. Data for three replicates for each of the Sequestin genes are presented in Figure S1G. (F) Phages engineered to express PF21825 genes overcome Thoeris defense. Shown are data for wild-type SBSphiJ phage, as well as SBSphiJ knocked-in with members of the PF21825 Sequestin family. Data represent PFUs per milliliter of phages infecting control cells (no system), or cells expressing the type I Thoeris system. Average of three biological replicates with individual data points overlaid.

Given the common fusions between Tad2 and Acb4, we hypothesized that new sponge proteins can be discovered based on their appearance fused to proteins from known sponge families. We therefore queried the Pfam database^21^ for protein families of unknown function that appear fused to either Tad2 (Pfam PF11195) or Acb4 proteins (PF13876). Tad2 was most commonly fused to proteins from Pfam family PF21825 (60 fusion instances). PF21825 was also the most common family fused to Acb4 proteins as well (91 fusion instances) (Fig. 1B, C).

PF21825 is a family of short (∼60–80 aa) proteins of unknown function predicted to adopt an alpha-helical structure (Fig. 1B, C). Most proteins with this family annotation appear in phage genomes or within prophages in bacterial genomes, suggesting that they primarily carry out a phage-specific function^21^. Despite frequent fusions with Tad2 and Acb4, proteins with the PF21825 domain most commonly appear as stand-alone proteins that do not contain other detectable domains except for PF21825^21^. Based on the preferential presence in phage genomes and frequent fusion with known sponges, we suspected that pfam PF21825 represents a previously uncharacterized family of viral sponges.

To test whether PF21825 proteins can inhibit bacterial immune signaling, we synthesized and cloned 13 members of this family and co-expressed each member with five distinct defense systems, each producing a different immune signaling molecule. These included type I Thoeris from *Bacillus cereus* MSX-D12 (producing 3′cADPR)^22^, type II Thoeris from *Bacillus amyloliquefaciens* Y2^22^ (His-ADPR), type I CBASS from *Escherichia albertii* MOD1-EC1698 (3′3′-cGAMP)^7^, Pycsar from *Escherichia coli* E831(cCMP)^10^, and Pycsar from *Bacillus sp*. G1 (speculated to produce cUMP).

We then challenged each strain with phages whose infection is naturally blocked by the respective defense system. Of the 13 PF21825 candidates tested, 10 affected the function of type I Thoeris, inhibiting it either partially or fully. This was observed both via infection in liquid cultures with low multiplicity of infection (MOI), where cultures expressing PF21825 proteins collapsed despite also expressing the Thoeris system (Fig. 1D, S1C-F), and via plaque assays, showing that co-expression of PF21825 proteins with the Thoeris system rendered the system inactive (Fig. 1E, S1G). To test if these proteins can inhibit defense also when expressed directly from the genome of the infecting phage, we engineered three of them into the genome of phage SBSphiJ under the control of the native promoter of the *tad1* sponge gene^11^. SBSphiJ, a phage naturally blocked by Thoeris, fully overcame this defense system when expressing either of the three PF21825 proteins (Fig. 1F). These results verify that PF21825 represents a family of phage-encoded Thoeris inhibitory proteins. None of the other tested defense systems were inhibited by the 13 members of the PF21825 family we experimented with (Fig. S1C-F).

The TIR domain protein of type I Thoeris (ThsB) produces 3′cADPR in response to phage infection. This signaling molecule activates the effector protein of the system (ThsA), which then cleaves NAD^+^ and depletes it from the infected cell. To determine whether PF21825 proteins interfere with 3′cADPR signaling, we incubated purified ThsA with lysates from phage-infected cells that express ThsB with or without proteins from the PF21825 family. As was previously shown^11,23^, lysates from infected cells expressing ThsB triggered the NADase activity of ThsA, confirming that the TIR protein ThsB produces 3′cADPR during infection by phage SBSphiJ (Fig. 2A). In contrast, lysates from infected cells co-expressing ThsB together with proteins from the PF21825 family failed to activate ThsA, suggesting that 3′cADPR signaling was impeded in these cells (Fig. 2A).

**Figure 2.**
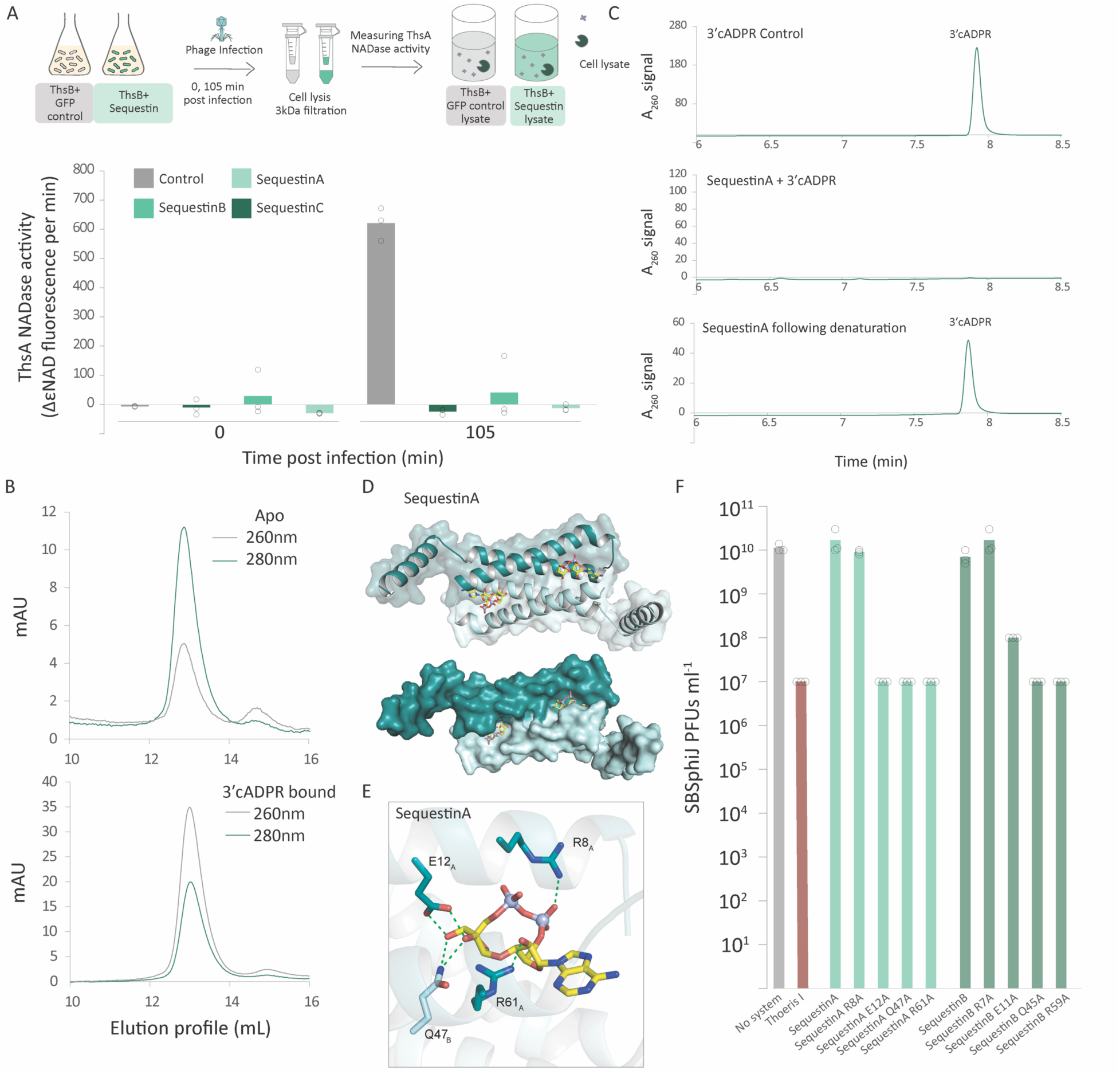
Sequestin is a family of viral sponges that bind and sequester 3ʹcADPR. (A) Sequestin proteins prevent accumulation of 3ʹcADPR in Thoeris-expressing infected cells. Cells expressing ThsB from *B. cereus* MSX-D12, or co-expressing both ThsB and genes from the Sequestin family, were infected with the phage SBSphiJ at a multiplicity of infection of 10. Cell lysates were extracted before infection (t=0) and 105 mins after infection, and were filtered to retain small molecules. Filtered lysates were then incubated with ThsA, and the NADase activity of ThsA was measured using a nicotinamide 1,N6-ethenoadenine dinucleotide (εNAD) cleavage fluorescence assay. Bars represent the mean of three experiments, with individual data points overlaid. Control represents experiments with lysates from cells expressing GFP together with ThsB. (B) Size-exclusion chromatography of SequestinA in apo state or following incubation with 3ʹcADPR. Light absorbance values at 260 nm and 280 nm are shown. (C) HPLC analysis of 3ʹcADPR incubated with either buffer (control, top lane) or with purified SequestinA (middle lane) at 1:1 ratio. Bottom lane shows the molecule release after the molecule-bound SequestinA was denatured by chloroform. (D) Structure of the AlphaFold3-predicted complex formed by a homodimer of SequestinA, together with 3ʹcADPR. Cartoon and surface representations are shown. (E) Close-up view of the predicted interactions of 3ʹcADPR with SequestinA from the model shown in panel E, highlighting key interacting residues. (F) Effect of point mutations in SequestinA and SequestinB on the ability of the sponge to cancel the defensive activity of type I Thoeris. Data represent PFU per milliter of SBSphiJ phage infecting negative control cells, cells expressing Thoeris, or cells co-expressing Thoeris and a sponge variant. Average of three replicates with individual data points overlaid.

To test whether PF21825 proteins may bind the signaling molecule of type I Thoeris, we purified a PF21825 protein, and performed size exclusion chromatography of this protein before and after incubation with 3′cADPR. This experiment revealed elevated 260nm/280nm light absorbance values for the protein pre-incubated with 3′cADPR, indicating that the protein directly binds the immune signaling nucleotide (Fig. 2B). Following incubation of the purified protein in a 1:1 ratio with 3′cADPR, the molecule was completely depleted from the solution, as confirmed by high-performance liquid chromatography (HPLC) (Fig. 2C). Furthermore, chloroform-denaturation of the 3′cADPR-bound protein released intact 3′cADPR molecules back into the medium (Fig. 2C). These results demonstrate that the studied protein is a viral sponge that binds and sequesters 3′cADPR to counteract Thoeris immunity.

Previously discovered families of viral sponge proteins were named after the defense systems that the founding member of the family was first shown to inhibit. Thus, as the first sponges to be discovered inhibited Thoeris defense, they were called Thoeris anti-defense 1 and 2 (Tad1 and Tad2)^11,18^. However, later studies showed that some members of the Tad1 and Tad2 sponge families can bind CBASS-produced signaling molecules^24^. To avoid ambiguous naming, we denote the PF21825 family of sponges with the name Sequestin. Since most Sequestin proteins we experimented with were derived from metagenomic databases of uncharacterized viruses, we refer here to metagenome-derived protein members of this family with the names SequestinA, SequestinB, and so on.

To gain deeper insight into the molecular mechanism by which Sequestin sponges sequester the immune signaling molecule, we modeled members of this sponge family using AlphaFold3 in the presence of 3′cADPR at various oligomeric states (Fig. 2D, S2). These proteins were predicted with high confidence to be folded as homo-dimers or homo-tetramers. In the homo-dimeric form, the two protomers are organized in an antiparallel manner. Two symmetrical, elongated pockets are formed in the interface between protomers, and AlphaFold3 places 3′cADPR in these pockets with high confidence (Fig. 2D, S2A). Despite substantial sequence divergence between Sequestin proteins modeled here, AlphaFold3 consistently places 3′cADPR in the same orientation within the Sequestin binding pockets, with high ipTM and pLDDT confidence scores, implying that this placement reflects natural binding.

We focused on protein SequestinA to further study the molecular interactions between the sponge and 3′cADPR. The model predicts that the placement of the immune signaling molecule in the pocket is coordinated via interactions with multiple conserved side chains. The adenine-proximal and adenine-distal ribose moieties are bound by R61 and E12 of protomer 1, respectively, and the short distance between the amino acid side chains and the hydroxyl groups of the respective riboses suggests hydrogen bonding (Fig. 2E). Q47 from the second protomer is also predicted to contribute to the interaction with the adenine-distal ribose (Fig. 2E). The phosphate groups are in close vicinity to the NH2 group of R8, suggesting that this residue coordinates the placement of the phosphates in the pocket. All residues indicated above are conserved in diverse members of the Sequestin sponge family and were predicted to interact with the signaling molecule also in other members of the Sequestin family. further supporting that they play a role in the interactions with 3′cADPR (Fig S2B). Substituting each of E12, Q47 or R61 into alanine resulted in loss of anti-defense activity, but an R8A substitution did not (Fig 2F). Mutations in the parallel residues in SequestinB, R7, E11, Q45 and R59A, resulted in the same phenotype (Fig 2F).

Using Foldseek^25^ and sequence-based analyses we detected 3,244 proteins belonging to the Sequestin sponge family in the IMG/VR v4 database of viral proteins^26^ (Table S1). Further analysis in a database of ∼25,000 fully sequenced phage genomes^27^ found these sponges in 639 phages, including phages infecting organisms belonging to 40 taxonomic genera from 6 bacterial phyla and one archaeal phylum (Table S1). Sequestin sponges were detected in the genomes of 14 phages from the BASEL collection^28^ all of which belonging to the Tevenvirinae group. Intriguingly, some well-studied model phages, including phages T2, T4 and T6, encode a protein from the Sequestin family. In phage T4 this protein is called Y16Q, a 64 amino acid long protein of unknown function. AlphaFold3 modeling showed that Y16Q folds as a homodimer with two pockets predicted to bind 3′cADPR, suggesting that this T4 protein can inhibit Thoeris similar to other members of the sponge family (Fig. S2A). Indeed, engineering Y16Q into phage SBSphiJ under the control of the *tad1* native promoter^18^ rendered the phage resistant to Thoeris, demonstrating that the phage T4 Y16Q protein is an anti-Thoeris sponge (Fig. 1F). Together, these results show that Sequestin is an anti-Thoeris sponge family abundant and widespread in phages in nature.

### Computational prediction of viral proteins that counter immune signaling

Examining the predicted structures of Sequestin sponges, as well as the determined crystal structures of other known sponges^11,18–20^, we noticed that they all share common biophysical properties. First, all sponges are encoded by short proteins (often less than 100 amino acids in size), and in all cases the ternary structure comprises a homo-oligomer: homo-dimers in the case of Sequestin, homo-tetramers in the case of Acb4 and Tad2, or homo-hexamers for Tad1 and Acb2. In all cases, the pockets that sequester the immune signaling molecules are formed at the interfaces between protomers, so that the final complex contains multiple binding sites for target molecules (up to 8 sites in the case of Tad1^24^). This unique protein complex organization allows phages to encode multi-molecule binding sponges while devoting little genomic space for this purpose.

Further analysis of crystal structures and AlphaFold predictions of known sponges in their apo forms showed that in almost all cases, the nucleotide binding pockets are deeply engraved or form large void spaces in the protein (Fig 3A). These spaces typically have an inner surface area of 200–700 Å^2^, and are almost always highly positively charged, likely to accommodate interactions with the negative charge of the phosphate groups that are present in all known nucleotide signaling molecules (Fig 3A). Given that known sponge proteins display the same biophysical traits despite belonging to evolutionarily unrelated families, we reasoned that new families of viral sponges can be discovered based on structure-guided analyses of large viral protein spaces. Under the premises of this hypothesis, short viral proteins that form homo-oligomers, in which multiple positively charged pockets are formed at the interface between protomers, can be candidate new sponges.

**Figure 3.**
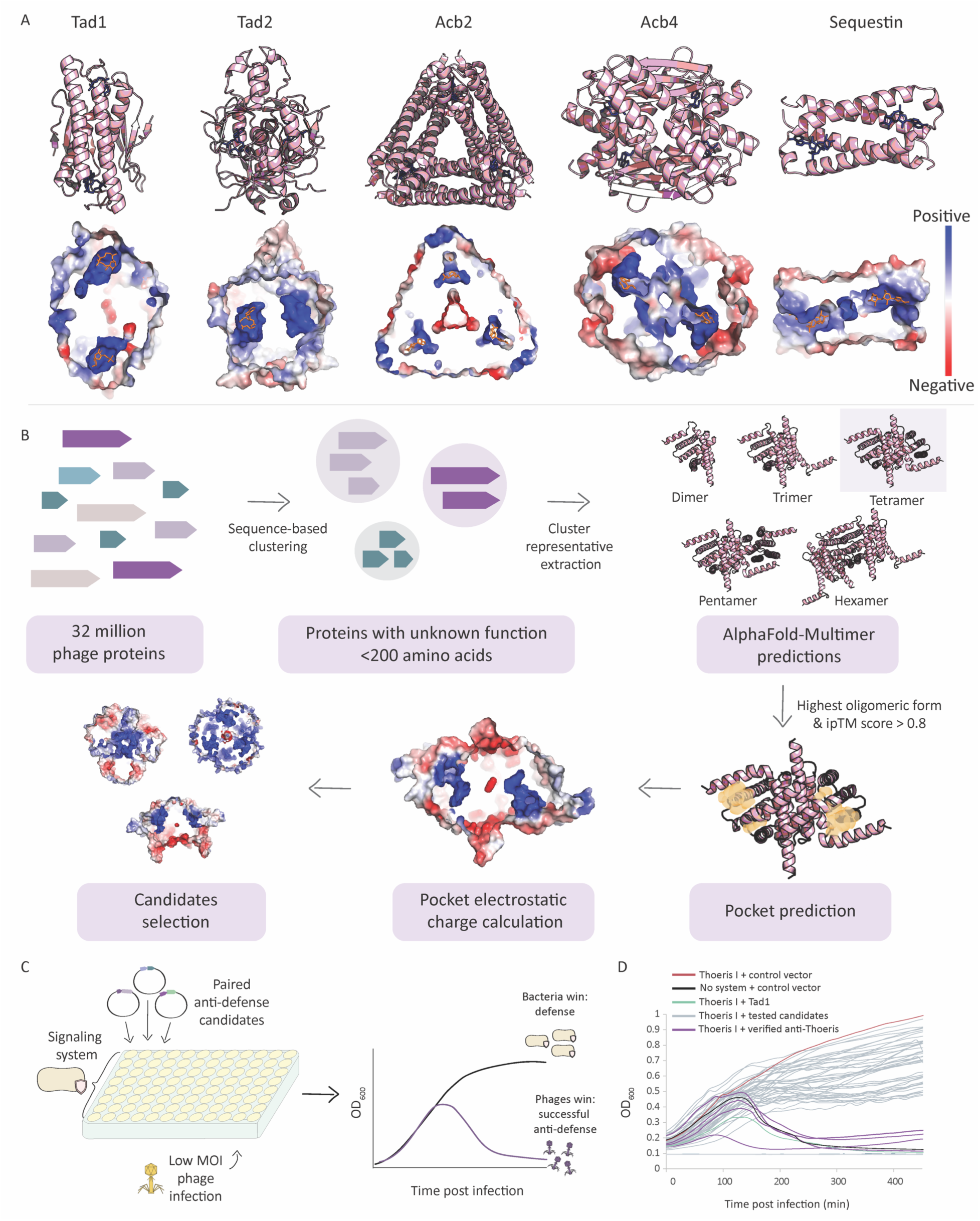
A structure-guided computational and experimental pipeline for the discovery of viral proteins that manipulate immune signaling. (A) Crystal structures of Tad1, Tad2, and Acb2 (PDB: 7UAW, 8SMG, 8IY2) and AlphaFold3-predicted structures for Acb4 (uniport accession: A0A0B3RSW4, ipTM = 0.93), and SequestinB (ipTM = 0.94), together with their respective ligands. Cross-sections into the proteins are presented, with surface as an electrostatic map in a blue-red scale. Blue blebs represent positively charged pockets within the proteins. A dimer unit of the Tad1 hexamer is presented for clarity. (B). A computational pipeline to predict viral proteins that interact with host immune signals. The phage protein space is clustered, and representative proteins are selected from clusters of short proteins of unknown function. High-scoring AlphaFold-Multimer predictions for homo-oligomers are analyzed by CastP^32^ and Autosite^33^ to identify pockets with sizes typical to those found in known sponge proteins. Proteins with positively charged pockets are further evaluated as possible new anti-defense proteins. (C) Schematic representation of the experimental set up. Liquid cultures of bacteria harboring the defense system are grown in a 96 well plate format. Each well is transformed with a different plasmid that expresses a pair of anti-defense candidates, with antibiotics added to select for cells that acquired the plasmid. Bacteria are subsequently infected with phages in a low MOI, and OD is measured to identify successful phage infections. (D) An example of an infection assay described in (C). Cells expressing the type I Thoeris system from *B. cereus* MSX-D12 were infected with phage SBSphiJ at MOI 0.01. Each curve represents growth of bacteria transformed with a pair of candidate anti-defense genes, with negative control cells transformed with a plasmid expressing GFP (black), and positive control cells transformed with a plasmid expressing Tad1 (green). Purple curves represent cases where the transformed plasmid allowed the phage to propagate and cause eventual culture collapse, suggesting that one of the genes in the transformed plasmid inhibited Thoeris defense.

To test this hypothesis we examined a dataset of roughly 32 million non-redundant proteins derived from approximately 2 million phage genome scaffolds found in the IMG/VR v3 database, as previously described (Methods)^29,30^. Proteins were clustered based on sequence similarity, excluding those with known functional annotations as well as those larger than 200 amino acids. A representative sequence from each cluster was analyzed using AlphaFold-Multimer^31^ to examine the likelihood of the protein to form a dimeric, trimeric, tetrameric, pentameric, or hexameric homo-oligomers. We then searched for pockets with internal surface areas sized 100–1000 Å^2^ within high-confidence homo-oligomeric predicted structures using the pocket prediction tools CastP and AutoSite^32,33^, and retained predicted structures in which multiple pockets were formed in the interfaces between protomers, and in which the internal surfaces of the pockets were positively charged (Methods) (Fig. 3B). Based on these parameters we prioritized a set of >120 candidate proteins for further experimentation. Although the selected proteins fulfill the theoretical structural constraints of sponges, we expect that only a minority of them represent real sponge proteins, as positively charged pockets can have roles in manipulation of non-immune nucleotides^34,35^ or other roles^36,37^.

We developed a high-throughput screening methodology to test the effect of 124 selected candidates on multiple defense systems. In this method, pairs of candidate sponges are transformed on a plasmid into bacterial strains in which a defense system was genomically integrated (Fig. 3C Methods). This procedure takes place in a 96-well plate format where candidate or control plasmids are transformed. The cultures are grown and passaged to fresh media under antibiotic selection to maintain the plasmid, and then the culture is subjected to phage infection in liquid media at low MOI. This approach identifies proteins that, when co-expressed with the defense system, cancel its activity and allow phage propagation and phage-mediated culture collapse (Fig. 3C). The benefit of this method is that it does not necessitate isolation of individual transformed colonies, which would have made the screen of >120 candidates vs multiple defense systems prohibitively labor intensive; the drawbacks are that only ∼80% of the attempted transformations were successful in the 96 well plate format, and that positive hits necessitate further verification (Table S2).

We used this method to transform 62 plasmids, each expressing a pair of candidate genes, to the five bacterial strains expressing the signaling defense systems type I Thoeris, type II Thoeris, CBASS, Pycsar (cCMP) and Pycsar (cUMP), and infected transformants with phages naturally restricted by the respective defense system (Fig. 3C,D). In cases where culture collapse was observed with at least one of the defense systems, the two genes in the pair were separated, and each gene was individually tested for its ability to inhibit the respective defense system using plaque assays (Table S2). Nine proteins were verified as exhibiting substantial anti-defense activity and were further investigated through biochemical assays (see below). Seven of them were found to directly interact with immune signaling molecules, and these were grouped into two distinct protein families that are further studied in detail below (Table S2).

### Lockin, a sponge family that inhibits types I and II Thoeris

Lockin is a large family of proteins predicted to fold into curved, elongated structures mostly comprising beta sheets (Fig S3A). AlphaFold-Multimer consistently predicts these proteins to fold into symmetrical, cog-like hexamers, with each monomer comprising a symmetrical segment of the cog (Fig. 4A,B). Positively charged pockets are formed between the segments, and these pockets are enclosed via a loop that emanates from one of the monomers (Fig. 4A,C). The surface areas of the predicted pockets range between ∼260–440 Å^2^, within the premises of the sizes of pockets in known sponges (Table S3). Mass photometry analysis of a purified protein verified that it purifies as a hexamer, confirming the oligomeric structure prediction of AlphaFold (Figure S3B).

**Figure 4.**
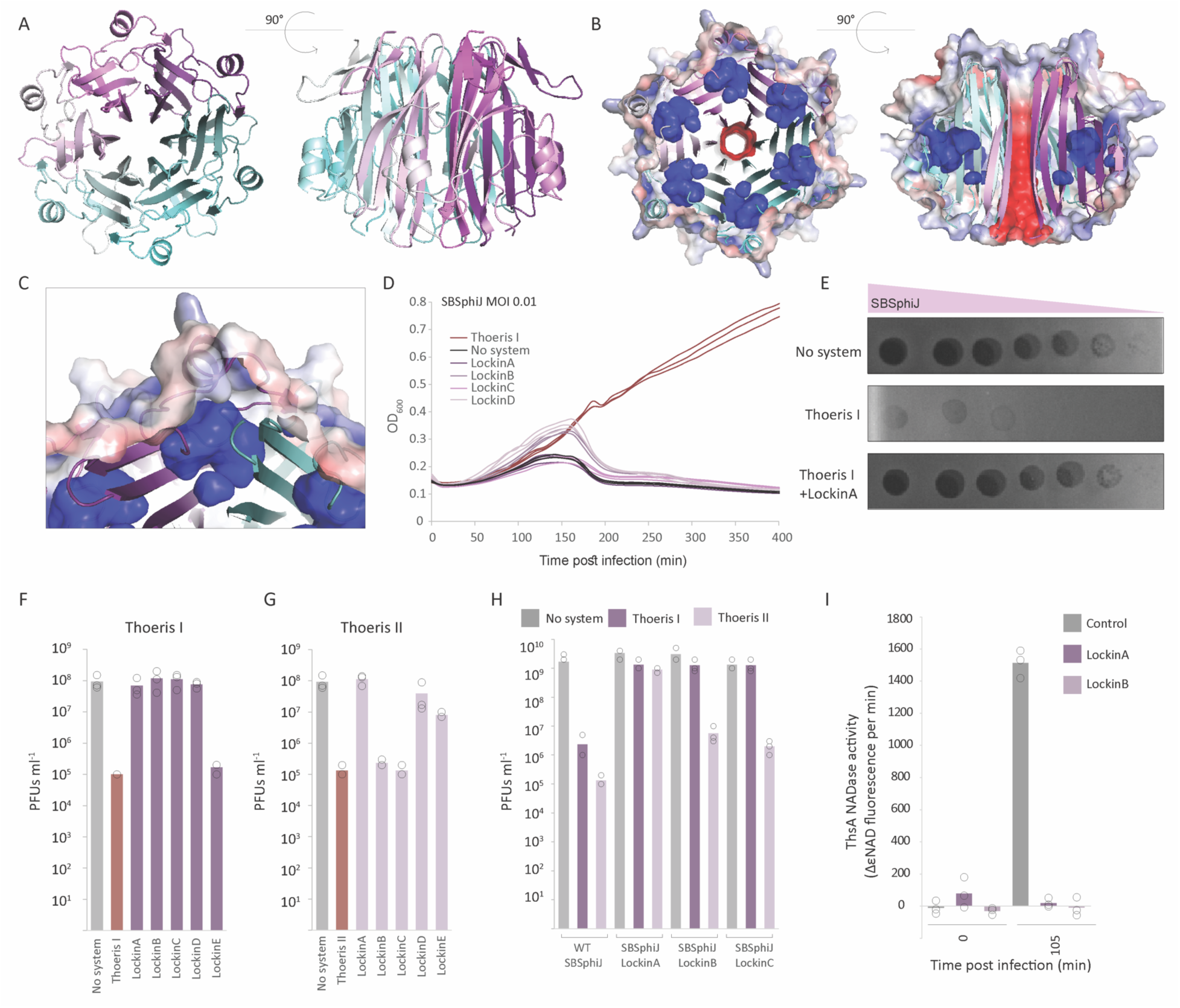
Lockin is a family of phage proteins that inhibit Thoeris defense. (A) Structure of the LockinA homo-hexamer, predicted by AlphaFold2-Multimer (ipTM = 0.78) in cartoon representation. (B) Cross sections in the LockinA structure model showing the void space pockets predicted to be formed within the protein. Top and side cross sections are shown on the left and right, respectively. Surface representation is displayed as an electrostatic map (blue-red scale for positive and negative charge, respectively). Blue blebs are positively charged pockets within the protein. (C) A close-up view on one of the positively charged pockets, showing that it is formed in the interface between protomers. (D) Growth curves of *B. subtilis* cells expressing type I Thoeris from *B cereus* MSX-D12 (red), or co-expressing type I Thoeris and a Lockin protein (shades of purple), or control cells expressing GFP instead of the Thoeris system (black). Cells were infected with SBSphiJ phage at an MOI 0.01. Results of three experiments are presented as individual curves. (E) A representative Lockin protein (LockinA) capable of overcoming type I Thoeris defense. Shown are tenfold serial dilution plaque assays, comparing the plating efficiency of phage SBSphiJ on bacteria that express the type I Thoeris alone or with LockinA. Images are representative of three replicates. (F) Plating efficiency of phage SBSphiJ on negative control cells, cells expressing type I Thoeris, and cells expressing both type I Thoeris and a Lockin gene. Data represent PFUs per milliliter, and bars show the average of three replicates with individual data points overlaid. (G) Same experiment as in panel E, but with type II Thoeris from *B. amyloliquefaciens* Y2. (H) Phages engineered to express Lockin genes overcome defense of types I and II Thoeris. Shown are data for wild-type SBSphiJ phage, as well as SBSphiJ knocked-in with members of the Lockin family. Data represent PFUs per milliliter of phages infecting control cells (no system), or cells expressing a type I or II Thoeris system. Average of three biological replicates with individual data points overlaid. (I) Lockin proteins prevent accumulation of 3ʹcADPR in Thoeris-expressing infected cells. Legend is as in Figure 2A.

Four Lockin proteins in our set exhibited strong activity against type I Thoeris (Fig. 4D-F), and two of them also inhibited type II Thoeris when co-expressed with the system (Fig. 4G). One additional Lockin protein exhibited activity against type II Thoeris exclusively (Fig. 4G). Engineering of Lockin proteins into phage SBSphiJ under the native promoter of *tad1* resulted in a phage resistant to one or both of the tested Thoeris systems (Fig. 4H), consistent with the co-expression results. Co-expression of Lockin proteins in phage-infected cells that express ThsB rendered the cells devoid of detectible 3′cADPR (Fig. 4I), similar to the results obtained for the anti-Thoeris sponge Sequestin above (Fig. 2A). These results verify that Lockin is a family of anti Thoeris proteins.

To further examine the interaction between Lockin and the signaling molecule, we purified the Lockin protein LockinA and incubated it with 3′cADPR. Size-exclusion chromatography followed by HPLC analysis showed an elevated 260nm/280nm absorbance ratio for the protein that had been pre-incubated with 3′cADPR, suggesting that it binds the signaling molecule (Fig. 5A). Indeed, 3′cADPR was eliminated from the solution after incubation with the Lockin protein, and reappeared following heat denaturation of the LockinA-3′cADPR complex (Fig. 5B). Together, these results verify that Lockin is a new family of anti-Thoeris sponges.

**Figure 5.**
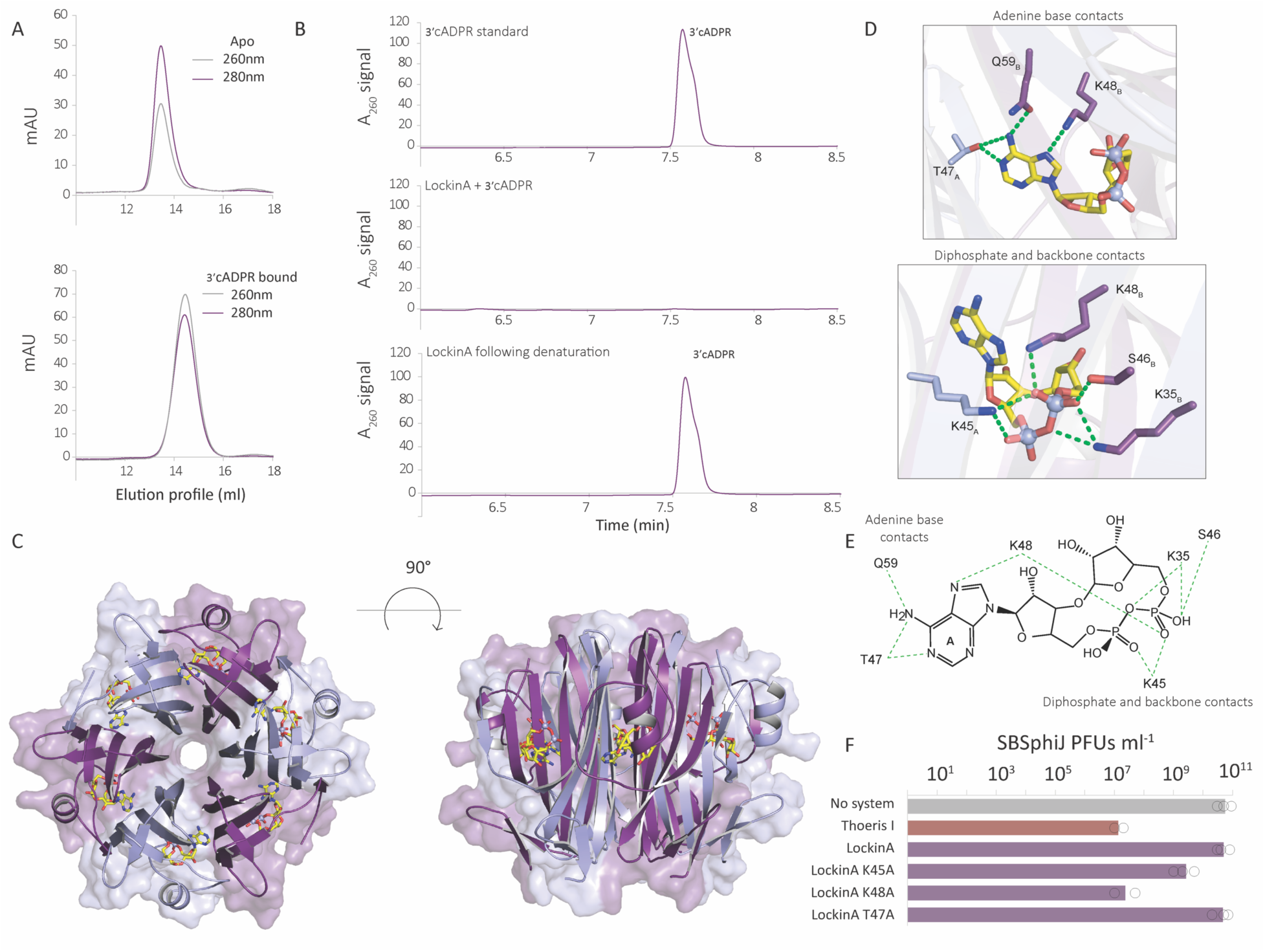
Lockin is a sponge protein that sequesters Thoeris signaling molecules. (A) Size-exclusion chromatography of apo state or 3ʹcADPR-bound LockinA. 3ʹcADPR-bound LockinA shows a substantial shift compared to LockinA in the apo state. (B) HPLC analysis of 3ʹcADPR incubated with either buffer (control, top lane) or with purified LockinA (middle lane) at 1:1 ratio. Bottom lane shows the molecule release after the molecule-bound Lockin was denatured in 98°C for 10 minutes. (C) Crystal structure of LockinA in complex with 3ʹcADPR. (D) Detailed view of LockinA residues that interact with the 3ʹcADPR adenine base and diphosphate backbone. Green dashed lines indicate hydrogen bonding interactions. Subscript denotes individual protomers within the Lockin hexamer. (E) A 2D map presenting a detailed view of 3ʹcADPR adenine base and diphosphate backbone and their LockinA residues interactions. (F) Effect of point mutations in LockinA on the ability of the sponge to cancel the defensive activity of type I Thoeris. Data represent PFU per milliter of SBSphiJ phage infecting negative control cells, cells expressing Thoeris, or cells co-expressing Thoeris and a sponge variant. Average of three replicates with individual data points overlaid.

To define the molecular basis of Lockin anti-Thoeris immune evasion, we determined a 1.6 Å X-ray crystal structure of Lockin in complex with 3′cADPR (Fig. 5C). Consistent with AlphaFold predictions, Lockin forms a hexameric assembly that sequesters six molecules of 3′cADPR (Fig. 5C). The 3′cADPR binding site resides in a deeply recessed pocket at the interface between each adjacent protomer (Lockin_A_ and Lockin_B_, respectively). The 3′cADPR ligand is coordinated by nucleobase-specific interactions between the adenine base and residue T47_A_ and Q59_B_ that read out the N1 and N6 positions of the nucleobase Watson-Crick edge (Fig. 5D,E). On the nucleobase Hoogsteen edge, Lockin residue K48_B_ makes further contacts with the nucleobase and coordinates the adenine N7 position (Fig. 5D,E). In addition to base-specific contacts, multiple Lockin residues stabilize the diphosphate backbone of 3′cADPR, including a network of polar contacts from K45_A_, K35_B_, S46_B_, and K48_B_ (Fig. 5D,E). Lockin point mutations in either K45 or K48 abolished or reduced the anti-defense activity of LockinA, as demonstrated by partial or complete restoration of immunity in cells co-expressing Thoeris and the mutated sponge (Fig. 5F). Mutating T47 did not change the anti-defense phenotype.

We detected 907 sequence and structural homologs of the verified Lockin proteins in the IMG/VR v4 database of viral proteins^26^ (Table S4). Analyzing homologs from isolate genomes, we found that Lockin proteins appeared in phages infecting bacteria from 11 taxonomic genera including *Bacillus*, *Rosebium*, *Pseudovibrio*, *Dehalobacter*, *Microcystis* and *Cohnella* (Table S4). In all cases, the proteins were annotated as phage proteins of unknown function, with no known domain annotation. We were not able to find members of this family within full-length sequenced and assembled phage genomes in the INPHARED database^27^, indicating that these proteins are under-represented in model phages whose genomes were fully sequenced.

### Acb5, a family of phage proteins that inhibit CBASS

Two phage proteins in our screen were able to inhibit the activity of the 3′3′-cGAMP-producing CBASS operon from *E. albertii* (Fig. 6A-C). Both these proteins, when co-expressed with the signal-producing CD-NTase enzyme of the *E. albertii* CBASS, prevented the accumulation of 3′3′-cGAMP in phage-infected cells (Fig. 6D). Despite low sequence similarity between the two proteins (Fig. S4A,B), both proteins are predicted to fold into structurally similar homo-tetrameric complex with high confidence AlphaFold-Multimer scores (Fig. 6E-F). Two large, positively charged pockets with internal surface areas of 425–530 Å^2^ are formed between loops at the interface between pairs of protomers (Fig. 6G). We denote this family of proteins Acb5 (Anti-CBASS 5).

**Figure 6.**
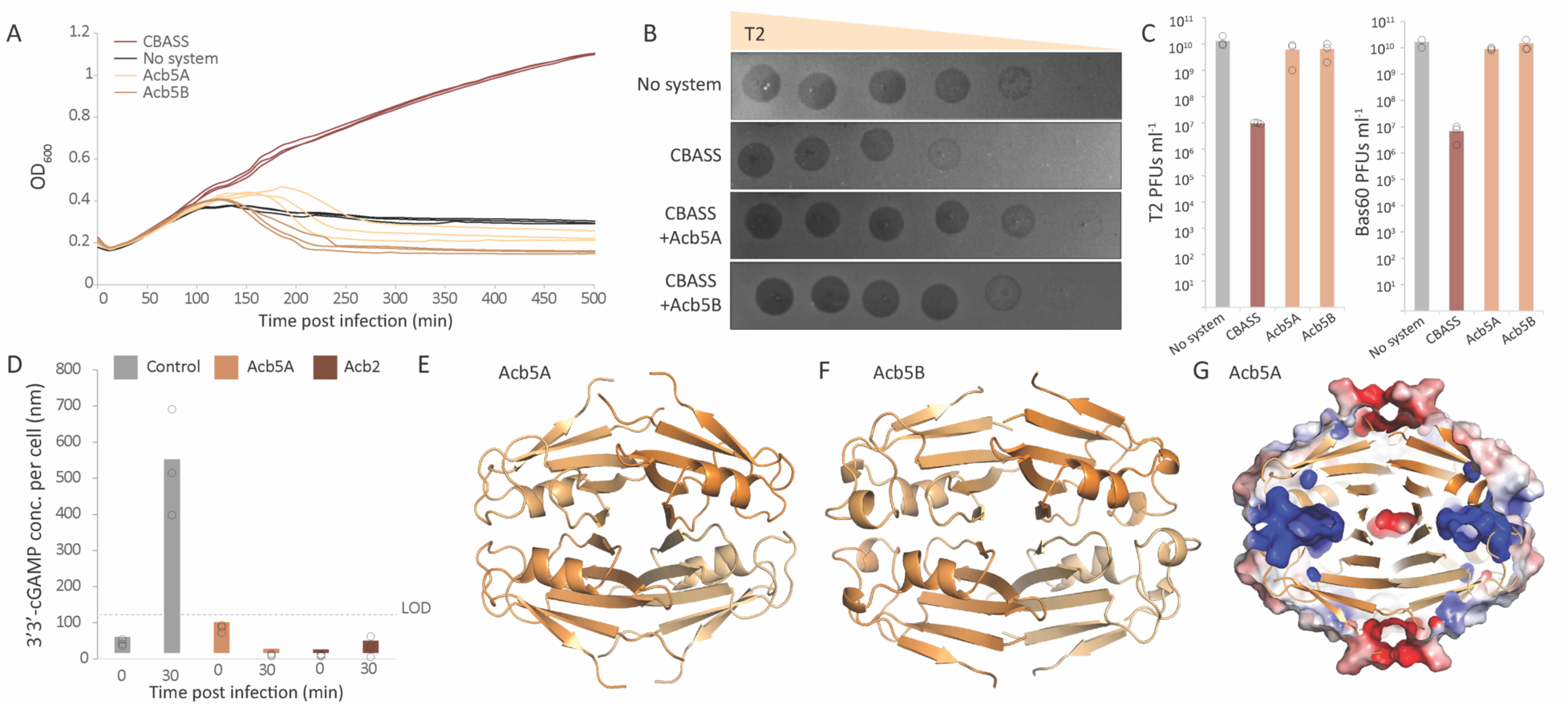
Acb5 proteins inhibit CBASS signaling. (A) Growth curves of *E. coli* cells expressing either a genomically-integrated CBASS system from *E. albertii* MOD1-EC1698 (red), co-expressing CBASS and Acb5 proteins (shades of orange), or negative control cells lacking the system (black), infected with T2 phage at an MOI 0.001. Results of three experiments are presented as individual curves. (B) Acb5 is capable of overcoming CBASS defense. Shown are tenfold serial dilution plaque assays, comparing the plating efficiency of T2 on bacteria that express the CBASS defense system, or bacteria co-expressing CBASS with Acb5 proteins, or a control strain that lacks the system. Images are representative of three replicates. (C) Plating efficiency of phage T2 and phage Bas60 on negative control cells, cells expressing CBASS, and cells expressing both CBASS and an Acb5 gene. Data represent PFUs per milliliter, and bars show the average of three replicates with individual data points overlaid. (D) Acb5 prevents cGAMP accumulation in infected cells. Cells expressing the CD-NTase enzyme of the *E. albertii* CBASS, or co-expressing the CD-NTase and Acb5A, were infected with the phage Bas60 at a multiplicity of infection of 10. Cell lysates were obtained before infection (0 mins) and during infection (30 min), and filtered to retain small molecules. cGAMP concentration in filtered lysates was measured using a 3ʹ3ʹ-cGAMP ELISA kit, and cGAMP concentration per cell was calculated based on estimated cell counts. Cells co-expressing CBASS and Acb2^19^ were used as positive control. Bars represent the mean of three experiments, with individual data points overlaid. Limit of detection is presented as a dashed line. (E-F) AlphaFold3-predicted tetramer complex for Acb5A (ipTM = 0.89) and Acb5B (ipTM=0.88). Protomers are presented in different shades of orange. (G) Cross section in the AlphaFold3-predicted tetramer complex for Acb5A with surface displayed as an electrostatic map (blue-red scale for positive-negative charges). Blue blebs represent positively charged pockets within the protein.

Surprisingly, thin-layer chromatography (TLC) analysis showed that incubation of purified Acb5 with radiolabeled 3′3′-cGAMP resulted in altered migration of the radiolabeled signal, leading us to suspect that Acb5 proteins may degrade cGAMP (Fig. 7A). To investigate this hypothesis, we incubated purified Acb5A with cGAMP and analyzed the resulting samples using HPLC. The analysis revealed that cGAMP was enzymatically digested into two distinct products (Fig. 7B,C). Further analysis using HPLC and mass spectrometry showed that the degradation produces had identical m/z values and retention times as standard 2′3′-cyclic AMP (cAMP) and 2′3′-cyclic GMP (Fig. 7C,D, S4C,D), suggesting that Acb5A cleaves cGAMP in two places and leaves 2′3′ cyclic phosphate groups on the cleavage products.

**Figure 7.**
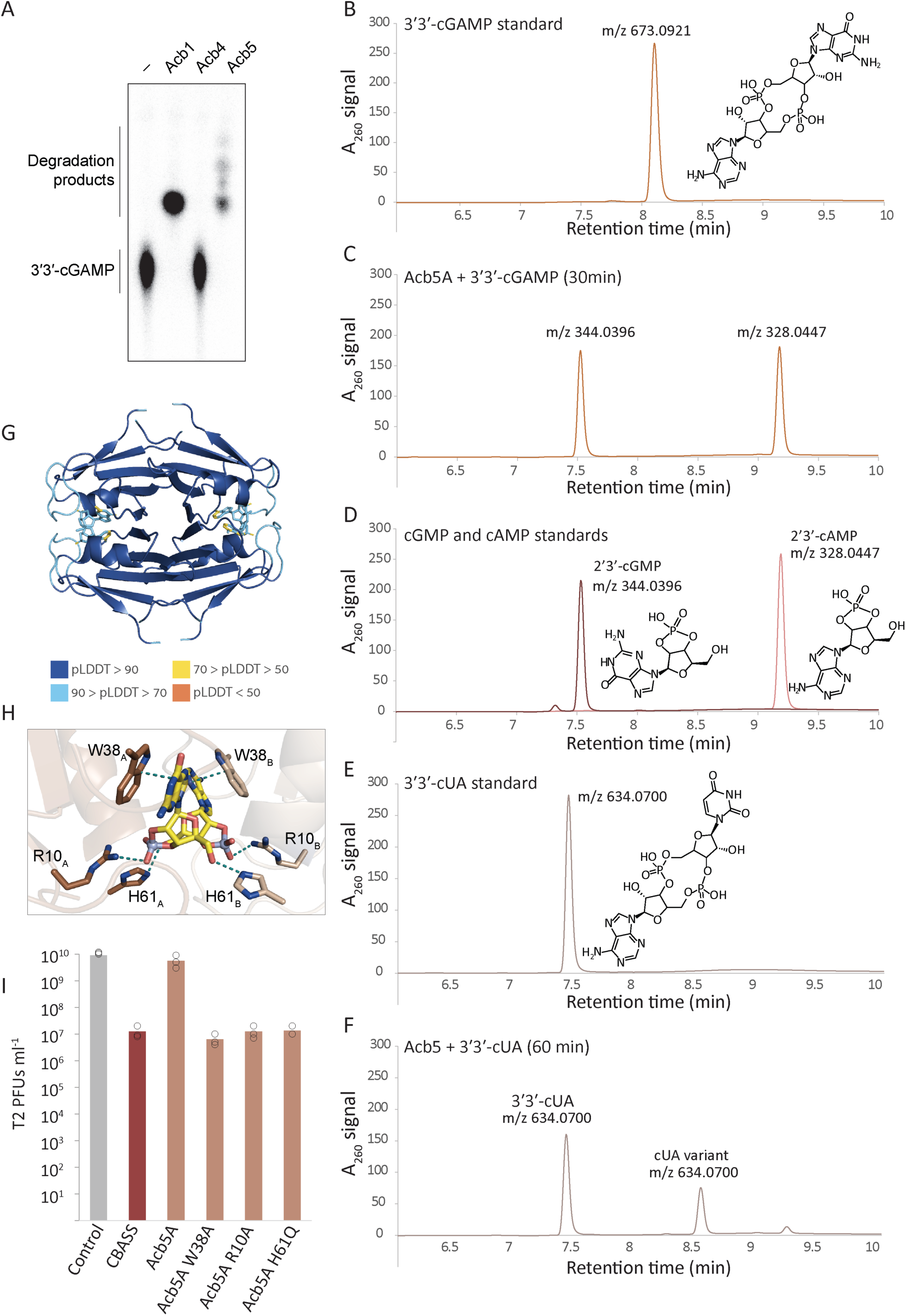
Acb5 is an enzyme that cleaves CBASS signaling molecules. (A) Thin-layer chromatography (TLC) analysis of Acb5A nuclease activity. Recombinant Acb5 was incubated with α^32^P-radiolabeled 3ʹ3ʹ-cGAMP, and degradation products were visualized by TLC. Purified Acb1 (an enzyme that cleaves 3ʹ3ʹ-cGAMP)^16^ and Acb4 (a 3ʹ3ʹ-cGAMP sponge)^20^ are included as controls. Data are representative of at least three independent experiments. (B) HPLC analysis of synthetic 3ʹ3ʹ-cGAMP. Y axis represents light absorbance at 260nm. (C) HPLC analysis of degradation products of 3ʹ3ʹ-cGAMP following 30 minutes incubation with purified Acb5A. M/Z values, as measured in mass spectrometry, are indicated for each product. Data are representative of 3 independent experiments. (D) HPLC analysis of synthetic 2ʹ3ʹ-cAMP and 2ʹ3ʹ-cGMP standards. (E) HPLC analysis of synthetic cUA. (F) Degradation products of 3ʹ3ʹ-cUA following 60 minutes incubation with purified Acb5A. (G) AlphaFold3-predicted tetramer complex for Acb5A with 2ʹ3ʹ-cAMP and 2ʹ3ʹ-cGMP (ipTM = 0.87, pTM=0.90), colors represent pLDDT scores. (H) Close-up view of the predicted interactions of 2ʹ3ʹ-cAMP and 2ʹ3ʹ-cGMP with Acb5A from the model shown in panel G, highlighting key interacting residues. (I) Effect of point mutations in Acb5A on the ability of the enzyme to cancel the defensive activity of CBASS. Data represent PFU per milliter of T2 phage infecting negative control cells, cells expressing CBASS, or cells co-expressing CBASS and an Acb5A variant. Average of three replicates with individual data points overlaid.

Incubation of Acb5A with 3′3′-cUA also resulted in partial degradation of cUA, suggesting that Acb5A can use 3′3′-cUA as a substrate, but to a lower efficiency than cGAMP (Fig 7E-F). HPLC and mass spectrometry analyses showed that the major degradation product has an m/z value identical to cUA but with a different retention time, suggesting that this product is the outcome of a single cleavage and phosphate cyclization event (Fig 7F). We hypothesize that cleavage takes place in the adenine moiety, implying a specificity of Acb5A for purines as substrates. These findings indicate that Acb5 is a family of enzymes that degrade CBASS-derived signaling molecules, thus inhibiting CBASS by disrupting immune signaling cascade.

AlphaFold3 consistently placed 3′3′-cGAMP, as well as the catalysis products 2′3′-cyclic AMP (cAMP) and 2′3′-cyclic GMP (cGMP), within the predicted pockets (Fig. 7G). The ligand-binding pocket is predicted to be symmetric and is formed by conserved residues from two adjacent protomers (Fig. 7H). The model predicts the conserved residue W38 from both protomers to interact with the adenine and guanine bases by pi-pi stacking. R10 and H61 are both predicted to interact with the phosphate moieties, and R10 is also placed in close proximity to the ribose moieties. Mutating either of these residues eliminated the anti-defense activity of Acb5A, supporting their predicted roles in the interaction with cGAMP (Fig. 7I).

We detected 1480 homologs of Acb5 proteins in the IMG/VR v4 database of viral proteins^26^ based on sequence and structural homology analyses (Table S5). Analyzing homologs from isolate genomes showed an extensive taxonomic diversity, with Acb5 proteins appearing in phages infecting both bacteria and archaea belonging to more than 20 taxonomic genera (Table S5). Acb5 proteins were especially abundant in phages infecting bacteria from the Actinobacteriota phylum, including *Streptomyces*, *Mycobacterium*, *Nonomuraea*, and many more (Table S5). Analysis of the INPHARED database of completely sequenced phage genomes^27^ found Acb5 homologs in 96 phage genomes, (Table S5). Consistent with the observations above, most of the Acb5 homologs in isolate phage genomes were found in phages infecting *Streptomyces* (66 phages) and *Mycobacteria* (14 phages). Homologs of Acb5 were also found in 11 phages infecting *Bacillus* species and 5 viruses infecting the Halophilic archeaeon *Haloarcula* (Table S5).

## Discussion

In this study, we showed how proteins with counter-immune activities can be discovered within large datasets of viral proteins based on the biophysical characteristics of predicted protein structural models. A major benefit of this approach is that it does not require *a priori* knowledge on the function of the studied proteins, nor it necessitates these proteins to be present in model organisms. An example for this is the discovery of Acb5, which is an anti-CBASS enzyme enriched in phages infecting *Streptomyces*, bacterial species not very commonly studied in the context of phage-bacteria interactions. Similarly, proteins from the Lockin family were not detected within sequenced genomes of model phages, demonstrating how unbiased structural analyses of phage proteomes can reveal anti defense proteins under-represented in known model viruses.

We find it remarkable that the well-studied phage T4 encodes multiple proteins that inhibit bacterial immune signaling systems. T4 was first found to encode anti-CBASS 1 (Acb1), an enzyme that cleaves and inactivates cyclic dinucleotides and cyclic trinucleotides^16^. Later studies showed that T4 also encodes anti-CBASS 2 (Acb2), a sponge that sequesters 3′3′-cGAMP^19^. Our study now shows that T4 also encodes a sponge from the Sequestin family, Y16Q, capable of inhibiting Thoeris defense. The observation that the same phage encodes multiple proteins that counteract diverse immune signaling systems attests to the importance of signaling-based systems in the phage-host arms race.

Through our unbiased approach, we also discovered Acb5 as the founding member of a new family of cGAMP-degrading enzymes, demonstrating further functional diversity of phage-encoded nucleases involved to evasion of host immunity. Notably, the Acb5 family does not share structural homology with Acb1 or any other immune evasion phosphodiesterase enzymes, indicating that phages have evolved multiple proteins for degrading host nucleotide immune signals^16,38,39^. In contrast to Acb1, which is a monomeric enzyme that hydrolyzes one bond in 3′3′-cGAMP and releases a linear product, Acb5 forms a two-fold symmetric active site that hydrolyzes both phosphodiester linkages in 3′3′-cGAMP and releases mononucleotide products. Together, these observations demonstrate that our unbiased systematic analysis of phage proteomes can be applied to uncover unexpected mechanisms that phages use to subvert host nucleotide immune signaling.

Although our approach is useful for the discovery of new sponges and nucleotide-cleaving counter immune enzymes, it cannot identify all phage sponges and enzymes. For example, Acb1 and Apyc1, enzymes that cleave and inactivate the immune signaling molecules cGAMP and cCMP/cUMP, respectively, do not include deep positively charged pockets^16^. Similarly, the internal surfaces of molecule-binding pockets in some members of known sponge families are not substantially positively charged. This was observed for example in the structure of the Acb4 sponge from phage SPO1, where the pocket residues in the sponge protein primarily interact with the nucleobases of the signaling molecules^20^. Different methods should be developed to discover viral sponges and enzymes that do not abide to the biophysical traits common to such proteins.

Most of the proteins we tested in our screens did not show anti-defense activities against the five defense systems included in this study. While many of these proteins likely represent false predictions, some of them might be true sponges that bind signaling molecules of defense systems not tested through our assays. The CBASS system we tested uses 3′3′-cGAMP as its signaling molecule, but other CBASS systems use different molecules including the cyclic dinucleotides 3′3′-cUA, 3′3′-c-di-UMP, 2′3′-cGAMP, 3′2′-cGAMP, the cyclic trinucleotides 3′3′3′-cAAG, 3′3′3′-cAAA and 2′3′3′-cAAA, and more^5,7–9^. Some of the non-verified candidate sponges might specifically bind one of these signaling molecules, a possibility that can be explored in future studies. Similarly, proteins in our study might inhibit type III CRISPR-Cas systems that produce cyclic oligo-adenylates as signaling molecules, or recently discovered defense systems that rely on inosine derivatives for immune signaling^40^.

Sponges emerge as a common strategy used by phages to inhibit bacterial immunity. Anti-Thoeris and anti-CBASS sponges were serendipitously detected in four separate unbiased studies aimed to understand how phages inhibit host immunity^11,18,19,41^, and our study now further demonstrates the abundance and diversity of sponges in phages. Despite the widespread utilization of sponges by viruses infecting bacteria, it is currently unknown whether viruses infecting eukaryotes use the same counter-defense strategy. Future application of our structure-guided sponge detection approach on proteomes of animal and plant viruses will enable unbiased searches for sponge proteins in the eukaryotic virus realm.

## Methods

### Detection of proteins fused to known sponge proteins

The online portal of the Pfam database^21^ was searched for Pfam protein families that appear fused to either Tad2 (Pfam PF11195) or Acb4 proteins (PF13876). Homologs of such proteins were taken from the online Pfam portal.

### Computational pipeline for prediction of positively charged pockets in phage proteome

∼32 million phage proteins from ∼2 million phage genome scaffolds were clustered based on sequence homology as previously described^29^, and ∼65,000 clusters of short proteins of unknown function with at least 20 members were further analyzed^29^. To generate multiple sequence alignment for the AlphaFold2 run, the databases of AlphaFold-Multimer 2.3.1^31^ with default parameters, with the addition of the IMG/VR v3 database to the default protein databases, were used. AlphaFold-Multimer was then used to predict the structure of a representative sequence^42^ from each cluster folded as a dimer, trimer, tetramer, pentamer and hexamer, with a single model predicted (model 1) for each homo-oligomer. For predicted structures with a model confidence score^31^ of at least 0.8, five models were calculated, and structures with an average model confidence score of at least 0.8 over the five models were retained for further analysis. For proteins with more than one possible oligomeric state (score > 0.8), the highest oligomeric form was chosen for further analysis.

Autosite^33^ and CastP^32^ with default parameters were ran on each retained homo-oligomeric structure to predict pockets within the structures. Structures in which CastP predicted at least one pocket with a surface area of >=100 Å^2^ and number of corner points >=150 were further examined, but ignoring pockets with more than 150 atoms. Structures in which Autosite predicted at least one pocket with a score >=60 were also further examined.

Electrostatic charge spatial maps of the proteins were generated using APBS ver. 3.1.1^43^ with the default parameters used in the PyMol APBS plugin. The discrete charge data obtained from the APBS analysis was linearly interpolated to calculate the charge value at the sampled points.

To calculate the surface charge of pockets predicted by the AutoSite software, the Autosite pocket output was first filtered to include only positions (“points”) where the distance to the closest atom in the protein was smaller than 1.2 Å. A charge positivity score was computed by calculating the fraction of points that have a charge greater than 5RT_ec_^−1^ out of the total number of points defining the pocket.

To calculate the surface charge of pockets predicted by CastP, the group of protein atoms that were predicted by CastP to participate in the pocket were analyzed. The solvent accessible surface was defined as the surface encompassing a radius of 1.4Å around each pocket atom, but excluding areas clashing with other atoms of the protein. Values for the van der Waals radius of atoms were the same values used in PyMOL. The solvent accessible surface was randomly sampled with uniform density of 0.48Å^-2^. A charge positivity score was computed by calculating the fraction of the sampled points that have a charge greater than 5RT_ec_^−1^.

For each protein the pocket with the highest positivity score in the Autosite analysis and the pocket with the highest positivity score in the CastP analysis were further considered. Proteins having at least one pocket with Autosite positivity score of 0.97, or at least one CastP-predicted pocket with positivity score of 0.3, were further examined as possible candidate scores. These thresholds were selected based on a set of 59 predicted structures spanning the structure space of Tad1, Tad2, and Acb2 for which the pockets were characterized previously^11,18,19^.

Proteins with pockets that passed the above-mentioned thresholds were searched for homologs in the IMG/VR v3 database using FoldSeek^25^, and were further examined manually. Candidates whose pockets were not conserved across protein homologs were discarded. Candidates having significant hhpred hit (e-value <= 0.05) to a protein of known function were also discarded.

### Bacterial strains and growth conditions

*E. coli* K-12 MG1655 and *B. subtilis* BEST7003 strains were grown in magnesium manganese broth (MMB; LB + 0.1 mM MnCl_2_ + 5 mM MgCl_2_) at 37 °C shaking at 200 rpm. Whenever applicable, the appropriate antibiotics were added at the following concentrations: For *B. subtilis* strains spectinomycin (100 μg ml^−1^) and chloramphenicol (5 μg ml^−1^), and for *E. coli* strains spectinomycin (50 μg ml^−1^) and kanamycin (50 μg ml^−1^).

Type I Thoeris from *Bacillus cereus* MSX-D12 (producing 3′cADPR), type II Thoeris from *Bacillus amyloliquefaciens* Y210 (His-ADPR), were cloned previously under their native promoters to the *amyE* locus of the *B. subtilis* BEST7003 genome^22^. Pycsar from *Bacillus* sp. G1 (cUMP) was cloned similarly under its native promoter to the *amyE* locus.

Type I CBASS from *Escherichia albertii* MOD1-EC1698 (3′3′-cGAMP)^7^, and Pycsar from *Escherichia coli* E831 (cCMP)^10^, were integrated into the genome of *E. coli* MG1655 using the Tn7 integration plasmid (spectinomycin resistance, induced by Anhydrotetracycline, Sigma cat.37919)^44^.

### Phage strains

The *B. subtilis* phage SBSphiJ (Genbank: LT960608.1) isolated by us in a previous study was used to perform all infection experiments for type I and type II Thoeris (MOI 0.01, unless stated otherwise). Phi105 (Genbank: NC_048631.1) was obtained from DSMZ (DSM HER46) and was used to infect Pycsar from *Bacillus* sp. G1 (MOI 0.01).

The *E. coli* T2 phage was obtained from DSMZ (16352) and phage BAS60 was obtained from the BASEL phage collection kindly contributed by Prof. Alexander Harms^28^. Both these phages were used to infect Type I CBASS from *E. albertii* (T2 MOI 0.001, Bas60 MOI 10). T5 was kindly contributed by Udi Qimron, and was used to infect Pycsar from *E. coli* E831 (MOI 0.01).

The phages were propagated on either *E. coli* MG1655 or *B. subtilis* BEST7003 by picking a single phage plaque into a liquid culture grown at 37 °C to an optical density at 600 nm (OD_600_) of 0.3 in MMB broth until culture collapse (or 3 hours in the case of no lysis). The culture was then centrifuged for 10 min at 3,200 *g* and the supernatant was filtered through a 0.2 µm filter to get rid of remaining bacteria and bacterial debris.

### Cloning and transformation of candidate anti-defense genes

Candidate anti-defense genes were codon optimized for *B. subtilis* and synthesized by Genscript Corp. as pairs under the control of the same IPTG promoter. The sequence ‘taataaggaggacaaac’ was added between the two open reading frames to generate a ribosome binding site. These constructs were subsequently cloned into the thrC-Phspank^11^ vector, which carries a low-copy pSC101s origin of replication and a kanamycin resistance marker. The resulting plasmid was then introduced into *Bacillus subtilis* BEST7003 cells, where the respective defense system was integrated into the amyE locus, or into *Escherichia coli* K-12 MG1655, where the defense system was integrated downstream of the gmlS gene under the control of a anhydrotetracycline-responsive promoter.

Transformations were performed using a 96-well format. Prior to transformation, *B. subtilis* BEST7003 cells with or without a defense system were cultured in MC medium at 37 °C for 3 hours, as described by Doron *et al*.^22^. MC medium was composed of 80 mM K2HPO4, 30 mM KH2PO4, 2% glucose, 30 mM trisodium citrate, 22 μg ml^−1^ ferric ammonium citrate, 0.1% casein hydrolysate (CAA) and 0.2% potassium glutamate. Aliquots of 200 µL were transferred to a deep 96-well plate, and 200 ng of the anti-defense candidate or control plasmid was added to each well. After a 3 hour incubation at 37 °C, 1 mL of MMB medium supplemented with spectinomycin (100 μg mL^−1^) and chloramphenicol (5 μg mL^−1^) was added to each well to dilute the MC medium, and the plate was incubated overnight at 30°C. On the following day, a 10 µL aliquot from each well was transferred into 990 µL of fresh MMB containing spectinomycin (100 μg mL^−1^) and chloramphenicol (5 μg mL^−1^) for a single passage. After overnight incubation at 37 °C, the cultures were used as starters for infection experiments in liquid cultures.

For transformation of *E. coli* K-12 MG1655, cells carrying the defense system integrated into their genome were grown in MMB medium until reaching an OD₆₀₀ of 0.3. The cells were then centrifuged at 3,200 × *g* for 5 minutes and resuspended in TSS medium at a 20X concentration^18^. A 50 µL aliquot of the cell suspension was distributed into a deep 96-well plate, and 50 ng of the anti-defense candidate or control plasmid was added to each well. The transformation protocol consisted of three incubation steps: 5 minutes on ice, 5 minutes at room temperature, and an additional 5 minutes on ice. Subsequently, 500 µL of antibiotic-free MMB was added to each well, followed by incubation with shaking at 37 °C for 1 hour. After this recovery period, 500 µL of MMB containing 2x antibiotics: spectinomycin (100 μg mL^−1^) and kanamycin (100 μg mL^−1^) was added to each well, to reach a final concentration of 50 μg mL^−1^ spectinomycin and 50 μg mL^−1^ kanamycin, and the plate was incubated overnight at 37 °C. On the next day, 10 µL from each well was transferred into 990 µL of fresh MMB containing spectinomycin (50 μg mL^−1^) and kanamycin (50 μg mL^−1^) for a single passage. After overnight incubation at 37 °C, the cultures were used as starters for infection experiments in liquid cultures.

### Infection in liquid culture

Overnight cultures of bacteria harboring the defense system, negative control lacking the system and system + candidate anti-defense gene pair were diluted 1:100 in MMB medium. For both *B. subtilis* and *E. coli* cultures, 1mM IPTG was added to induce expression of candidate anti-defense proteins. For *E. coli* cultures anhydrotetracycline (aTC, Sigma cat.37919) was added to induce the expression of the defense system (800nM and 40nM for CBASS and Pycsar, respectively). Cells were incubated at 37 °C while shaking at 200 rpm until early log phase (OD₆₀₀ of 0.3), and infected in a 96-well plate containing 20 μL of phage lysate. Plates were incubated at 30°C with shaking in a TECAN Infinite200 plate reader and OD₆₀₀ was followed with measurement every 10 min.

### Plaque assays

Phage titer was determined using the small drop plaque assay method^45^. A 300 μl of overnight culture of *E. coli* or *B. subtilis* was mixed with 30 ml MMB 0.5% agar supplemented with 1mM IPTG (as well as 800nM and 40nM of aTC for CBASS and Pycsar, respectively) and poured into a 10-cm square plate followed by incubation for 1 h at room temperature. Tenfold serial dilutions in MMB were carried out for each of the tested phages and 10 µL drops were put on the bacterial layer. After the drops had dried up, the plates were inverted and incubated overnight at room temperature (for systems expressed in *B. subtilis)* or 37 °C (for systems expressed in *E. coli*). PFUs were determined by counting the derived plaques after overnight incubation and lysate titer was determined by calculating PFUs per milliliter. When no individual plaques could be identified, a faint lysis zone across the drop area was considered to be 10 plaques. The efficiency of plating was measured by comparing plaque assay results for control bacteria and those for bacteria containing the defense system and/or the defense system with a candidate anti-defense gene.

### Preparation of filtered cell lysates for NADase enzymatic assay

For generating filtered cell lysates, *B. subtilis* BEST7003 cells co-expressing the candidate anti-defense proteins and the Thoeris system from *B. cereus* MSX-D12 in which ThsA was inactivated (ThsB + ThsAN112A) were used as described previously^23^. The genes SequestinA, SequestinB, SequestinC, LockinA and LockinB were integrated in the thrC locus and expressed from an inducible Phspank promoter^11^. The Thoeris system was integrated in the amyE locus and expressed from its native promoter. Controls included cells expressing only the ThsB + ThsA(N112A) Thoeris system with GFP. All cultures were grown overnight and then diluted 1:100 in 250 ml MMB medium supplemented with 1 mM IPTG and grown at 37 °C, 200 rpm shaking for 120 min followed by incubation and shaking at 25 °C, 200 rpm until reaching an OD_600_ of 0.3. At this point, a sample of 50 ml was taken as the uninfected sample (time: 0 min), and the SBSphiJ phage was added to the remaining culture at an MOI of 10. Flasks were incubated at 25 °C with shaking (200 rpm), for the duration of the experiment. Samples of 50 ml were collected 105 minutes post infection. Immediately after sample removal the sample tubes were centrifuged at 4 °C for 10 min at 3,200 *g* to pellet the cells. The supernatant was discarded, and the pellet was flash frozen and stored at −80 °C. To extract cell metabolites from frozen pellets, 600 μL of 100 mM Na phosphate buffer (pH 8.0) was added to each pellet. Samples were transferred to FastPrep Lysing Matrix B in a 2 ml tube (MP Biomedicals, cat #116911100) and lysed at 4 °C using a FastPrep bead beater for 2 × 40 s at 6 ms^−1^. Tubes were then centrifuged at 4 °C for 10 min at 15,000 *g*. The supernatant was then transferred to an Amicon Ultra-0.5 Centrifugal Filter Unit 3 kDa (Merck Millipore, no. UFC500396) and centrifuged for 45 min at 4 °C, 12,000 *g*. Filtered lysates were taken for *in vitro* ThsA-based NADase activity assay.

### ThsA NADase enzymatic assay

The ThsA protein was expressed and purified as previously described^18^, and ThsA-based NADase activity assay for the detection of 3′cADPR was carried out as described previously^23^. NADase reaction was carried out in black 96-well half-area plates (Corning, cat #3694). In each reaction microwell, purified ThsA protein was added to cell lysate, or to 100 mM sodium phosphate buffer pH 8.0. A 5 μL volume of 5 mM nicotinamide 1,N6-ethenoadenine dinucleotide (εNAD^+^, Sigma, cat #N2630) solution was added to each well immediately before the beginning of measurements, resulting in a final concentration of 100 nM ThsA protein in a 50 µL final volume reaction. Plates were incubated inside a Tecan Infinite M200 plate reader at 25 °C, and measurements were taken at 300 nm excitation wavelength and 410 nm emission wavelength. The reaction rate was calculated from the linear part of the initial reaction.

### Preparation of filtered cell lysates for cGAMP measurement

Cells expressing type I CBASS from *E.* albertii MOD1-EC1698 lacking the effector (delta TM) were used, co-expressing the candidate anti-defense protein Acb5A, a positive control of Acb2 or a negative control GFP. Cultures were grown overnight and then diluted 1:100 in 200 ml MMB medium supplemented with 1 mM IPTG and 800 nM aTC and grown at 37 °C, 200 rpm until reaching an OD_600_ of 0.3. At this point, a sample of 50 ml was taken as the uninfected (time 0 min) sample, and the Bas60 phage was added to the remaining culture at an MOI of 10. Flasks were incubated at 37 °C with shaking (200 rpm), for the duration of the experiment. Samples of 50 ml were collected 30 minutes post infection. Immediately after sample removal the sample tubes were centrifuged at 4 °C for 10 min at 3,200 *g* to pellet the cells. The supernatant was discarded, and the pellet was flash frozen and stored at −80 °C. To extract cell metabolites from frozen pellets, 600 μL of 100 mM Na phosphate buffer (pH 8.0) was added to each pellet. Samples were transferred to FastPrep Lysing Matrix B in a 2-ml tube (MP Biomedicals, number 116911100) and lysed at 4 °C using a FastPrep bead beater for 2 × 40 s at 6 m s^−1^. Tubes were then centrifuged at 4 °C for 10 min at 15,000g. The supernatant was then transferred to an Amicon Ultra-0.5 Centrifugal Filter Unit 3 kDa (Merck Millipore, no. UFC500396) and centrifuged for 45 min at 4 °C, 12,000 *g*. The amount of cGAMP in filtered lysates was measured using a 3′3′-cGAMP ELISA kit (Arbor Assays cat. K073-H) according to manufacturer’s instructions. Limit of detection was calculated during the experiment using a cGAMP standard curve, according to manufacturer’s instructions.

### Synthesis of radiolabeled cyclic nucleotide signals

Cyclic nucleotides were synthesized using the following purified recombinant enzymes: *Vibrio cholerae* DncV^9^: 3′3′-c-di-AMP, 3′3′-cGAMP, and 3′3′-c-di-GMP; and *Yersinia aleksiciae* CdnE: 3′3′-cUA^7^. Cyclic nucleotide synthesis was performed at 37 °C for ∼20 hours in 10 μL reactions containing a final concentration of 5 µM recombinant enzyme, 20 μM appropriate nucleotide triphosphates (NTPs), trace amounts of appropriate α-^32^P-labeled NTP (Revvity), 25 mM KCl, 5 mM MgCl_2_, 1 mM MnCl_2_, 1 mM TCEP and 50 mM Tris-HCl pH 7.5 (DncV) or pH 9.0 (*Ya*CdnE). Following overnight incubation, reactions were centrifuged at 17,000 × *g* for 1 minute to remove any precipitated protein. To digest unincorporated NTPs, clarified synthesis reactions were treated with 2 μL of Quick CIP (NEB) followed by incubation for ∼30 minutes at 37 °C and heat inactivation for 2 minutes at 80°C. To confirm cyclic nucleotide synthesis, ∼1 μL of the final reaction product was spotted on a PEI cellulose thin-layer chromatography plate (Sigma Aldrich) and developed in a 1.5 M KH_2_PO_4_ (pH 3.8) buffer for 90 minutes. Plates were dried at room temperature for ∼20 minutes, exposed to a storage phosphor screen, and imaged with a Typhoon Trio Variable Mode Imager System (GE Healthcare).

### Analysis of cyclic nucleotide degradation by thin-layer chromatography

Thin-layer chromatography was used to analyze Acb5A cyclic nucleotide degradation. Degradation reactions were performed at 37 °C in 5 μL reactions containing 5 μM recombinant Acb5, 5 μM of α-^32^P-labeled cyclic dinucleotide, 50 mM Tris-HCl pH 7.5, 5 mM MgCl_2_, 10 mM KCl, and 1 mM TCEP. Following 30 minutes incubation, reactions were boiled at 95 °C for 3 minutes and centrifuged at 17,000 × *g* for 1 minute. 1 μL of the boiled reaction was spotted on a 20 cm × 20 cm PEI cellulose thin-layer chromatography plate (Sigma Aldrich) and developed in a 1.5 M KH_2_PO_4_ (pH 3.8) buffer for 90 minutes. Plates were dried at room temperature for ∼20 minutes, exposed to a storage phosphor screen, and imaged with a Typhoon Trio Variable Mode Imager System (GE Healthcare).

### Lockin–3′cADPR crystallization and structure determination

Crystals of the Lockin–3′cADPR complex were grown using hanging-drop vapor diffusion method for ∼12 days at 18 °C. Recombinant Lockin was diluted to 10 mg mL^−1^ in a buffer containing 20 mM Tris-HCl pH 7.5, 250 mM KCl, and 1 mM TCEP. The diluted protein was incubated with 500 μM 3′cADPR (Biolog Life Science Institute) on ice for ∼30 minutes. The resultant protein mixture was allowed to equilibrate to 18 °C and crystals were grown in 96-well trays containing 70 μL reservoir solution and 800 nL drops. Drops were mixed 1:1 with purified protein and reservoir solution (11.5% PEG-6000, 0.1 M HEPES pH 7.5, and 5% (v/v) MPD). Crystals were cryo-protected with reservoir solution supplemented with 20% (v/v) glycerol and 500 μM 3′cADPR (Biolog Life Science Institute) and harvested by flash-freezing in liquid nitrogen. X-ray diffraction data were collected at Advanced Photon Source (beamline 24-ID-E), and data were processed on the RAPDv2 platform using XDS^46^. Experimental phase information was determined by molecular replacement using Phaser-MR in PHENIX^47^ and a model of Lockin generated using AlphaFold3^48^. Model building was performed using Coot^49^ and refined using PHENIX. A summary of crystallographic statistics is provided in Table S6. All structural figures were generated using PyMOL (Version 2.5.4, Schrödinger, LLC).

### Protein expression and purification

The anti-defense genes were cloned (TWIST Bioscience, USA) into the expression vector pET28-His-bdSumo. pET28-His-bdSumo was constructed by transferring the His14-bdSUMO cassette from the K151 expression vector generously donated by Prof. Dirk Görlich from the Max-Planck-Institute, Göttingen, Germany^50^ into the expression vector pET28-TevH^51^.

Each protein was expressed in *E. coli* LOBSTR-BL21(DE3)-RIL (Kerafast, USA) by induction with 500 μM IPTG at 16 °C overnight. The culture was harvested and lysed on ice by a EmulsiFlex-C3 (Avestin, Canada) in a lysis buffer (20 mM HEPES pH 7.3, 0.4 M NaCl, 1 mM DTT, 30 mM imidazole). After centrifugation (20,000 *g*, 30 min, 4 °C), the supernatant was filtered through a 0.4 µm filter, and sample was loaded on a HisTrap FF 5 mL column (Cytiva) prewashed with lysis-buffer. The column was subsequently washed with lysis-buffer followed by a wash buffer containing high salt (20 mM HEPES pH 7.3, 1 M NaCl, 1 mM DTT, 30 mM imidazole). The protein was eluted using elution buffer (buffer −20 mM HEPES pH 7.3, 0.4 M NaCl, 1 mM DTT, 300 mM imidazole), and was inserted into a 3.5 MWCO 15 mL slide-A lyzer^TM^ (Thermoscientific) with 0.4 mg of His-bdSENP1 protease in order to cleave the target protein from its His-bdSUMO tag and dialyzed against cleavage buffer (20 mM HEPES, 125 mM KCl, 1mM DTT) overnight at 4°C. The sample was centrifuged again (20,000 *g*, 30 min, 4°C), filtered, supplemented with NaCl and imidazole to final conc. of 275mM NaCl and 40mM imidazole. The sample was then reapplied to a HisTrap FF 5 mL column (Cytiva) in order to recapture the His-bdSUMO tag uncleaved precursor and the His-tagged bdSENP1 protease. Flowthrough containing the cleaved anti-defense protein was then passed through an Amicon ULTR-15 30 MWCO filter, in order to remove any remaining residual cleaved His-SUMO tag and His-tagged bdSENP1 protease. The protein was then stored at −80 °C.

### Mass photometry

Mass photometry^52^ measurements were acquired using a OneMP mass photometer (Refeyn Ltd). Microscope coverslips (no. 1.5, 24 × 50 Marienfeld, 0107222) were cleaned by sequential sonication in 50% isopropanol (HPLC grade) in Milli-Q H2O, and Milli-Q H2O (5 min each), followed by drying with a clean nitrogen stream. Four reusable culture-well gaskets 3 mm diameter × 1 mm depth (Sigma, GBL103250-10EA) were cut, washed with ethanol followed by MilliQ water and dried using nitrogen stream. Dried gaskets were placed at the center of the coverslip, where each well was used for one measurement. To establish focus, fresh 100 mM sodium phosphate (pH 8) buffer (15 µL) was first loaded into the well, the focal position was identified and secured in place for the entire measurement, with an autofocus system based on total internal reflection. Molecular weights were calibrated using pure BSA and bovine IgGs standards.

Purified LockinA protein was diluted to a concentration of 10nM. For each measurement, 5 µL of protein was introduced into the measurement well with the loaded buffer and mixed several times. After autofocus stabilization, movies were recorded for a duration of 120 s. Data acquisition was performed using AcquireMP (Refeyn Ltd, v.2.3) and data analysis was performed using DiscoverMP (Refeyn Ltd, v.2.3).

### Knock-in of Sequestin and Lockin candidates into phage SBSphiJ

The DNA sequence of the genes was amplified from the respective pSG-thrC-Phspank plasmid, or in the case of T4-Y16Q from the phage genome itself, using KAPA HiFi HotStart ReadyMix (Roche, cat. #KK2601) with the following primer pairs:

LockinA: ttttatacctataggaggtagctATGAAAGAAAAAGATCTTGGAATCACTGA,

ttttccctccagttgttcTTAAATTTTTCCGATAAAACGAGTGTTGC.

LockinB: ttttatacctataggaggtagctATGACCGTGGAGAAAACATTGAC,

ttttccctccagttgttcTCACGGATCGATAAGATCTACGTAGC.

LockinC: ttttatacctataggaggtagctATGAACATGGACAAAAAGGAAGAGAA,

ttttccctccagttgttcTTACGGGCGTGTGATTGAATATGTG.

SequestinA: tcaaattttatacctataggaggtagctatgGAGAAGTCTGTTTTTGATCGCCTGC, tttgccaagtgttttccctccagttgttcttaTTCATTAATCGTAGATTTCGCCAACGC.

SequestinB: tcaaattttatacctataggaggtagctatgCAAGACTTTCAACAGCGGGTA,

tttgccaagtgttttccctccagttgttcttaGATCTCAAAGCCCGCAATGCG.

T4-Y16Q: tcaaattttatacctataggaggtagctatgttagcttatcaagcacgagtaaaag,

tttgccaagtgttttccctccagttgttcttacttgaattgtgcaattcttttctctagac.

The backbone fragment with the upstream and downstream genomic arms (±1.2 kb) for the integration site of candidate was amplified from the plasmid used previously for knock-in of the *tad1* gene^11^ (F-gaacaactggagggaaaacacttg, R-agctacctcctataggtataaaatttg). Cloning was carried out using the NEBuilder HiFi DNA Assembly cloning kit (NEB, cat # E5520S) and the cloned vector was transformed into NEB 5-alpha competent cells. The cloned vector was subsequently transformed into the thrC site of *B. subtilis* BEST7003. To amplify phages, the anti-defense-containing *B. subtilis* BEST7003 strain was infected with phage SBSphiJ with a multiplicity of infection of 0.1 and cell lysate was collected. Phages with knock-in were selected by Cas13a-gRNA-SBSphiJ strain^53^. The selection was made on an agar plate in the presence of 0.2% xylose to induce Cas13. Resistant phages were purified three times on *B. subtilis* BEST7003. Purified phages were verified again for the presence of the anti-defense candidate using PCR and sequencing.

### Distribution of anti-defense proteins in phage genomes

Homologs of verified anti-defense genes were identified in the IMG/VR v4 database by conducting sequence-based and structure-based homology searches as described previously^29^. For this, 5.5 million phage scaffolds labeled as ‘‘high-confidence virus’’ were downloaded from the IMG/VR v4 database. Homologous sequences of the anti-defense proteins detected in this study were identified using the ‘‘search’’ option of MMseqs2 release 13-45111 with the parameters **‘**-c 0.8 –cov-mode 2’’. To identify structural homologs, the downloaded proteins from IMG/VR v4 were clustered using the ‘‘cluster’’ option of MMseqs2 with default parameters. Next, a representative sequence was extracted from each cluster containing at least 30 non-identical members, and its structure was predicted using AlphaFold2 version 2.3 with default parameters, resulting in 182,179 phage protein structures. Structures of the anti-defense proteins were searched against this set of 182,179 phage protein structures using Foldseek release 5.53465f065 with default parameters. Hits with probability of 1.0 and no additional protein domains outside of the homology region were collected with all their cluster members as structural homologs of the anti-defense proteins. Protein members belonging to clusters containing an additional domain outside of the homology region were defined as homologs if they also had any significant sequence similarity to the verified anti-defense proteins, using the “search” option of MMseqs2 with default parameters. Homologs of anti-defense proteins were detected in the INPHARED database and in the BASEL phage collection using the ‘‘search’’ option of MMseqs2 with the parameter **‘**-c 0.8’’, using all of the anti-defense homologs detected in IMG/VR v4 as queries.

### SequestinA and LockinA *in vitro* sponge experiment

SequestinA and LockinA were expressed and purified as described above in the protein expression and purification section. Sponge and ligand (3′cADPR, Biolog cat. C 404) were mixed in a 1:1 ratio to a final concentration of 100 uM, in 100 mM sodium phosphate buffer pH 8.0. Ligand and sponge, as well as a control containing the ligand only, were incubated for 10 minutes at room temperature. Following the incubation the samples were filtered through a 3 kDa MWCO filter, and flow through was collected and analyzed by HPLC. The upper fractions (containing the protein + ligand complexes) were washed by successive concentration and dilution in a 3 kDa MWCO filter with five 1:20 dilutions in PBS. For LockinA the upper fraction was then boiled at 98 °C for 10 min, pelleted at 13,500 *g* for 20 min, and filtered through a 3 kDa MWCO filter. The filtrate was collected and assessed by HPLC. For SequestinA, the upper fraction was then subjected to chloroform denaturation as follows. 200 µL of the sample were mixed with 200 µL of phenol:chloroform:isoamyl alcohol (25:24:1), vortexed for 1 min and centrifuged at 10 000 *g* for 10 min. The aquatic phase was taken and mixed with 200 µL chloroform:Isoamyl alcohol (24:1), vortexed for 1 min, centrifuged at 10,000 *g* for 10 min and 10ul of the aquatic phase was taken and analyzed by HPLC.

### Acb5A enzymatic reaction

Acb5A enzymatic reaction was carried out by mixing a purified Acb5A with its ligand (3′3′-cGAMP, biolog cat. C117) in a 1:100 ratio (1 µM Acb5A, 100 µM 3′3′-cGAMP) in 0.1 M phosphate buffer (sodium phosphate pH8). 200 µL were collected at each time point. Time point 0’ was taken before adding the enzyme, and time point 30’ was taken following 30 minutes of incubation at 30°C. Similarly, Acb5A was mixed with cUA (Biolog cat. C357) in a ratio of 1:100 (1 µM Acb5A, 100 µM cUA) in 0.1M phosphate buffer. Time point 0’ was taken before adding the enzyme, and time point 60’ was taken following 60 minutes of incubation at 30°C. Following incubation, the samples were filtered through a 3 kDa MWCO filter, and flow through was collected and analyzed by HPLC.

### HPLC analysis

10 µL of the samples were analyzed using HPLC. HPLC of the obtained fraction was performed using Agilent 1260 and chromatography SUPELCOSIL™ LC-18-T HPLC Column. The following protocol was used for all runs: 1 min of mobile phase A 100%, 2 min 75% A and 25% B, 2 min 50% A and 50% B, 2 min 20% A and 80% B and 3 min 100 % A, 1 mL/min flow rate. Mobile phase A was 20mM potassium phosphate pH 6 and B was 20mM potassium phosphate pH 6 in 20% methanol.

### LC-MS polar metabolite analysis

The sample collected for HPLC were also analyzed by UPLC–HRMS without dilution. The analyses were carried out on a Waters SYNAPT-XS Q-Tof mass spectrometer (Manchester, UK) with an electrospray ionization (ESI) source. The spectra cGAMP, cGMP, cAMP and cUA were recorded in the negative ion mode within a mass range from 100 to 1200 m/z. The parameters were set as follows: capillary voltage at 1.5 kV, cone gas flow at 50 l/h, source temperature at 140°C, and cone voltage at 20 V. The desolvation temperature was set at 600°C, and the desolvation gas (N_2_) flow rate was set at 800 l/h. Lock spray was acquired with Leucine Encephalin (m/z=554.2615 in negative mode) at a concentration of 200 ng/ml and a flow rate of 10 μl/min once every 10 s for 1 s period to ensure mass accuracy. Waters MassLynx v4.2 software was used for data acquisition and data processing. The analytes were separated using Waters Premier Acquity UPLC system. The gradient elution was achieved with Waters Acquity Premier HSS T3 Column, 1.8 μm, 2.1 × 100 mm at 0.25 ml/min flow rate, 35°C. Mobile phase A consisted of 0.1% of aqueous formic acid (Fisher Scientific A117-50) and mobile phase B consisted of 0.1% formic acid in acetonitrile:water (95:5). Identification of substrates and products was done by MS and retention time, and validated by the injection of a commercially available standard.

### Analytical SEC analysis of apo and ligand-bound SequestinA and LockinA

17.2 µL volume of LockinA (218 µM) or 15 µL of SequestinA (250 µM) protein was incubated with 7.5 µL of 3′cADPR (5mM) and mixed with PBS buffer (final volume of 100 µL) at 25°C for 10 minutes. As a control, the apo protein was incubated under identical conditions without the ligand. Following incubation, each sample was loaded onto a Superdex 200 Increase 10/300 GL (Cytiva) SEC column, pre-equilibrated with PBS buffer. Chromatography was performed at a flow rate of 0.6 mL/min at 4°C using an ÄKTA purifier, and sample was monitored by absorbance at both 260 nm and 280 nm.

### Prediction of protein-ligand contacts

AlphaFold3^48^ was used to generate structural models for all sponges or enzymes with their ligands. RING 4.0^54^ was applied to identify possible ligand-interacting residues together with manual examination for evolutionary conserved residues adjacent to the ligand.

## Data and materials availability

Coordinates and structure factors of the LockinA–3′cADPR complex have been deposited in the Protein Data Bank under the accession number 9P8L. All other data from the manuscript are available in the manuscript or in supplementary materials.

## Acknowledgements

We thank members of the Sorek and Kranzusch lab for constructive discussion during this study; Y. Peleg and S. Albeck for assistance with protein expression R.S. was supported, in part, by the European Research Council (grant ERC-AdG GA 101018520), the Israel Science Foundation (MAPATS grant 2720/22), the Minerva Foundation with funding from the Federal German Ministry for Education and Research, a research grant from the Estate of Hermine Miller, the Institute for Environmental Sustainability (IES) and the Center for Immunotherapy at the Weizmann Institute of Science, and the Knell Family Center for Microbiology. The research was supported in part by grants to P.J.K. from the Burroughs Wellcome Fund PATH program, The G. Harold and Leila Y. Mathers Charitable Foundation, the Cancer Research Institute, the Parker Institute for Cancer Immunotherapy, the Massachusetts Consortium on Pathogen Readiness (MassCPR), the Gates Foundation (INV-083469), and the National Institutes of Health (1DP2GM146250-01). R.B.C. is supported through a Landry Cancer Biology Research Fellowship (Harvard Faculty of Arts and Sciences). X-ray data were collected at the Northeastern Collaborative Access Team beamlines, which are funded by the National Institute of General Medical Sciences from the National Institutes of Health (P30 GM124165). The Eiger 16M detector on the 24-ID-E beam line is funded by a NIH-ORIP HEI grant (S10OD021527). This research used resources of the Advanced Photon Source, a U.S. Department of Energy (DOE) Office of Science User Facility operated for the DOE Office of Science by Argonne National Laboratory under Contract No. DE-AC02-06CH11357. This publication resulted from the data collected using the beamtime obtained through NECAT BAG proposal # 311950.

## Supplementary Figures

**Figure S1.**
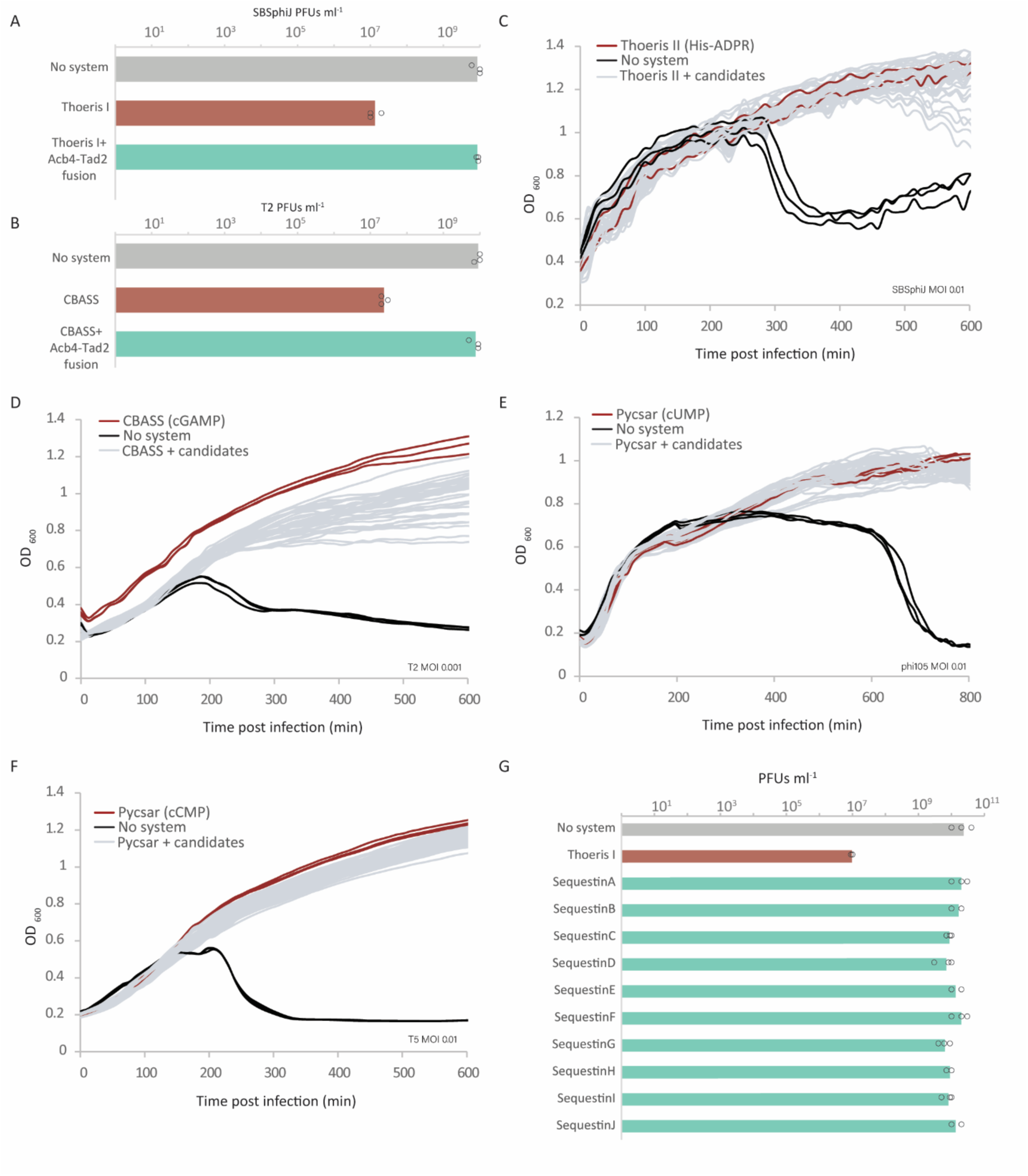
Sequestin proteins inhibit type I Thoeris. (A) Plating efficiency of phage SBSphiJ on control cells, cells expressing type I Thoeris from *Bacillus cereus* MSX-D12, and cells co-expressing the system with a fusion protein of Acb4 and Tad2 from *Pasteurella* phage Pm86. Results from plaque assay data in PFUs per milliliter are presented. Bars represent the average of three replicates with individual data points overlaid. (B) Plating efficiency of phage T2 on control cells, cells expressing CBASS from *Escherichia albertii* MOD1-EC1698, and cells co-expressing the system with the Acb4 and Tad2 fusion protein from *Pasteurella* phage Pm86. Results from plaque assay data in PFUs per milliliter are presented. Bars represent the average of three replicates with individual data points overlaid. (C-F) Sequestin proteins do not inhibit other defense systems tested in this study. Growth curves of *B. subtilis* cells expressing either type II Thoeris (C), CBASS (D), Pycsar (cUMP) (E) and Pycsar (cCMP) (F). Strains co-expressing the defense system with a Sequestin gene (grey), or expressing the defense system alone (red) were infected at a low MOI of SBSphiJ, T2, T5, phi105, for type II Thoeris, CBASS, Pycsar (cCMP) and Pycsar (cUMP), respectively. Results of three experiments are presented as individual curves. In each panel, negative control cells express GFP instead of the defense system (black). (G) Plating efficiency of phage SBSphiJ on control cells, cells expressing type I Thoeris, and strains coexpressing both type I Thoeris and a Sequestin protein. Results from plaque assay data in PFUs per milliliter are presented. Bars represent the average of three replicates with individual data points overlaid.

**Figure S2.**
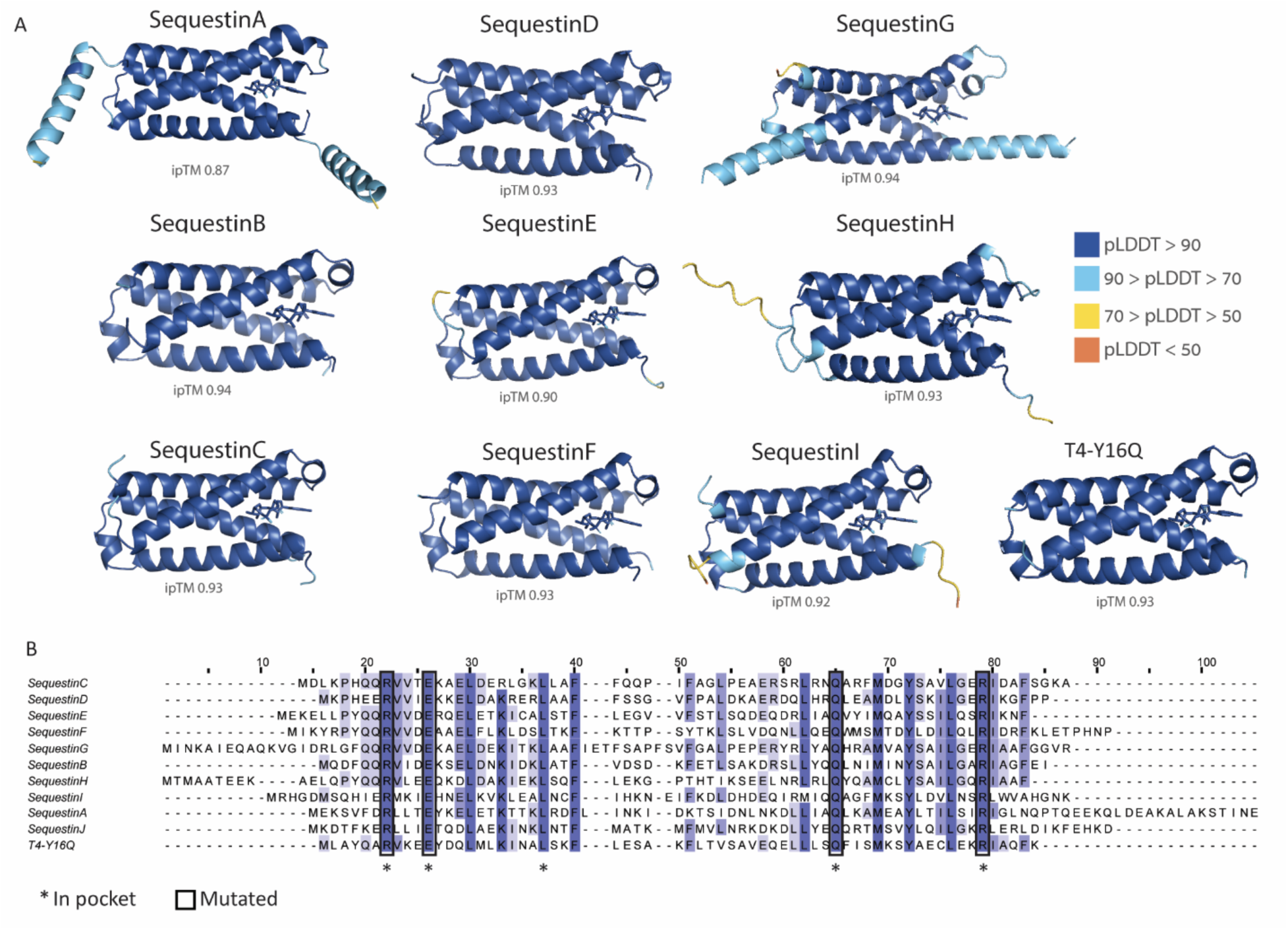
Sequestin proteins fold as homo-dimers with pockets accommodating 3ʹcADPR. (A) AlphaFold3-predicted homo-dimer complexes for the Sequestin proteins that showed anti-defense phenotypes in this study, co-folded with 3ʹcADPR. Colors represent pLDDT scores. (B) Sequence alignment of Sequestin proteins. Conserved residues are in blue. Asterisks indicate residues that are conserved and found in the pocket. Squares indicate residues that were mutated in this study.

**Figure S3.**
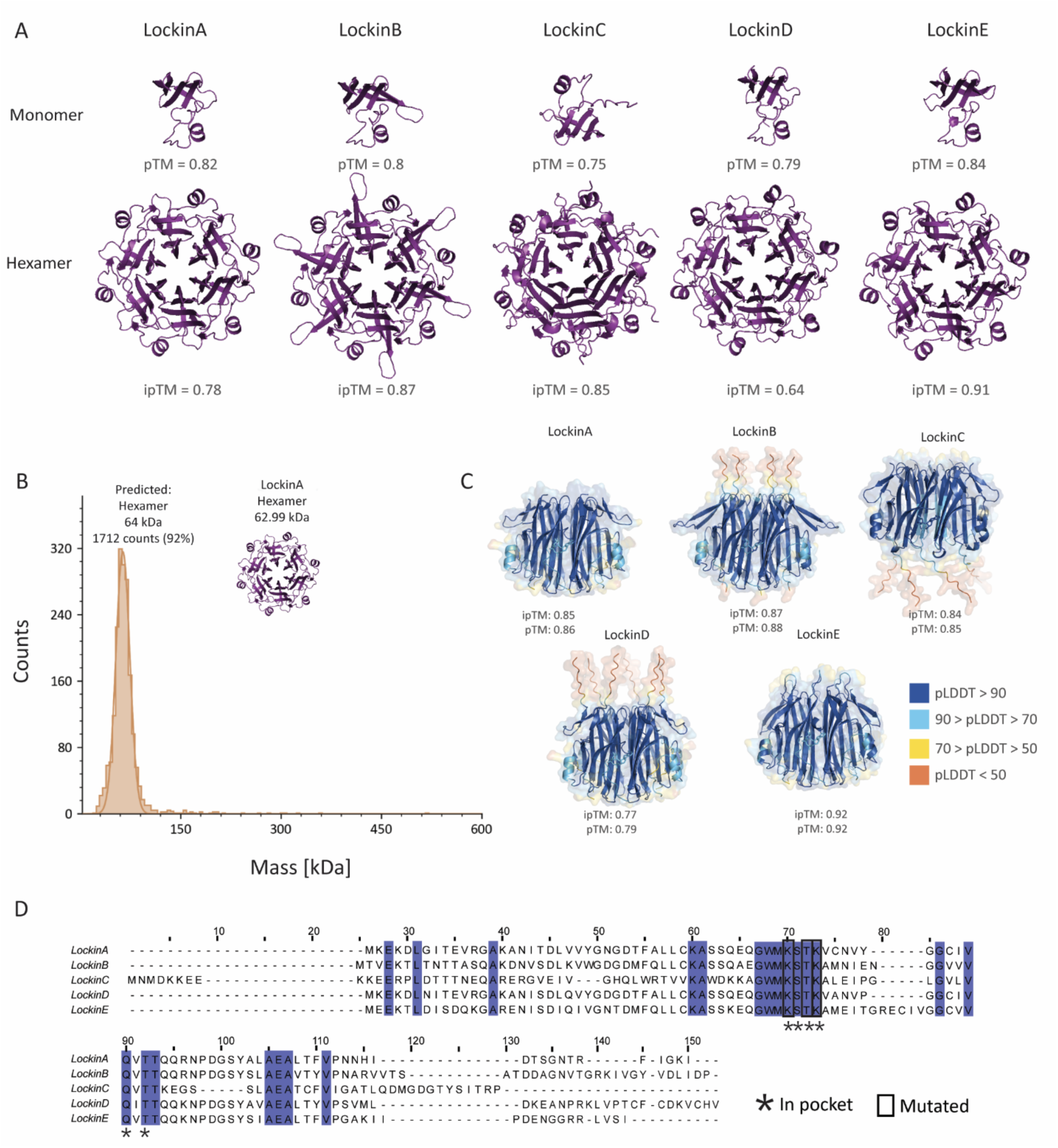
Lockin proteins form hexameric complexes with conserved pockets. (A) AlphaFold3-predicted monomer and homo-hexamers for Lockin that were shown experimentally in this study to inhibit Thoeris defense. (B) Histogram showing the masses of individual purified LockinA protein particles. Protein masses in solution were measured using mass photometry. (C) AlphaFold3-predicted homo-hexamer complexes for the Lockin proteins that showed anti-defense phenotypes, co-folded with 3ʹcADPR. Colors represent pLDDT scores. (D) Sequence alignment of Lockin proteins. Conserved residues are in blue. Asterisks indicate residues that are conserved and found in the pocket. Squares indicate residues that were mutated in this study.

**Figure S4.**
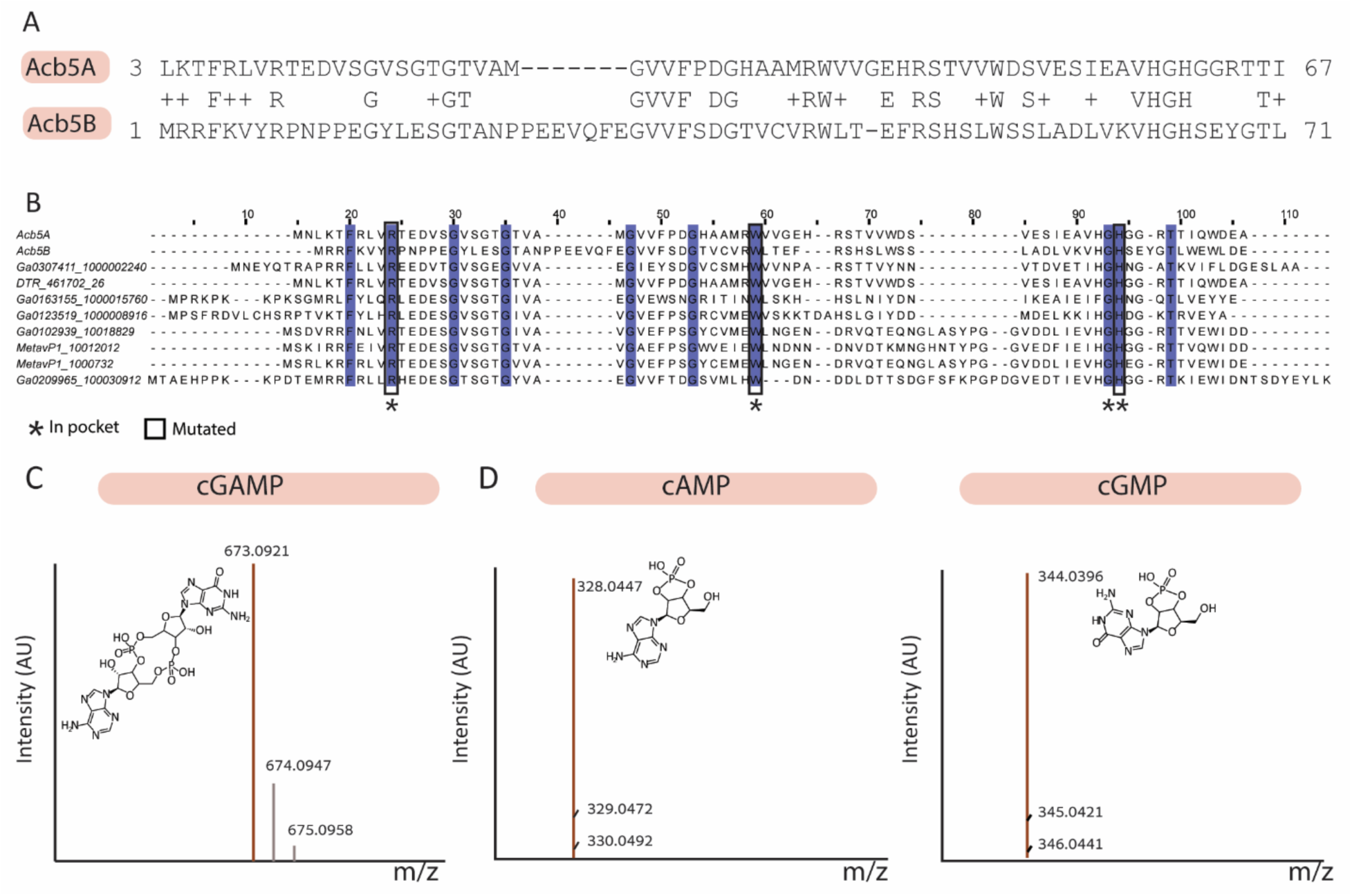
Acb5 proteins are enzymes that cleave cGAMP. (A) Sequence alignment of Acb5A and Acb5B, both were found to inhibit CBASS defense. (B) Sequence alignment of Acb5 proteins shown to inhibit defense (Acb5A, Acb5B) together with 8 additional randomly selected homologs. Conserved residues are in blue. Asterisks indicate residues that are conserved and found in the pocket. Squares indicate residues that were mutated in this study. (C-D). Liquid chromatography coupled with mass spectrometry (LC-MS) analysis. (C) LC-MS of 3ʹ3ʹ-cGAMP standard before incubation with Acb5A. (D) LC-MS detection of 2ʹ3ʹ-cAMP and 2ʹ3ʹ-cGMP following 30 minutes of incubation of Acb5A with 3ʹ3ʹ-cGAMP. Masses of detected products and molecule composition are indicated.

## Supplementary tables

Table S1. Sequestin homologs in the IMG/VR, INPHARED and BASEL databases

Table S2. Candidate inhibitors of signaling systems

Table S3. Pocket dimensions of known anti defense sponges

Table S4. Lockin homologs in the IMG/VR database

Table S5. Acb5 homologs in the IMG/VR and INPHARED databases

Table S6. A summary of crystallographic statistics of LockinA

