## Supplementary material for "Structural modeling reveals viral proteins that manipulate host immune signaling": Table S6

**Table S6. Crystallographic Statistics, Related to Figures 5**

|  | **Lockin–3′ADPR** |
| --- | --- |
| Resolution (Å)^a^ | 80.33–1.62 (1.65–1.62) |
| Wavelength (Å) | 0.97920 |
| Space group | P 2 2_1_ 2_1_ |
| Unit cell: a, b, c (Å) | 46.58, 74.42, 80.33 |
| Unit cell: α, β, γ (°) | 90.00, 90.00, 90.00 |
| Molecules per ASU | 3 |
| Total reflections | 472,935 (23,633) |
| Unique reflections | 36,306 (1,774) |
| Completeness (%)^a^ | 100 (100) |
| Multiplicity^a^ | 13.0 (13.3) |
| *I/σI^a^* | 13.5 (1.6) |
| CC(1/2)^b^ (%)^a^ | 99.8 (82.3) |
| Rpim^c^ (%)^a^ | 3.7 (45.2) |
| Resolution (Å) | 80.33–1.62 |
| Free reflections | 1,818 |
| R-factor / R-free | 17.66 / 20.93 |
| Bond distance (RMS Å) | 0.005 |
| Bond angles (RMS °) | 0.840 |
| No. atoms: protein | 2,042 |
| No. atoms: ligand / ion | 105 |
| No. atoms: water | 370 |
| Average B-factor: protein | 24.82 |
| Average B-factor: water | 36.47 |
| Ramachandran plot: favored | 100% |
| Ramachandran plot: allowed | 0.00% |
| Ramachandran plot: outliers | 0.00% |
| Rotamer outliers | 0.45% |
| MolProbity^d^ score | 1.10 |
| Protein Data Bank ID | 9P8L |

^a^ Highest resolution shell values in parentheses

^b^ (Karplus and Diederichs, 2012)

^c^ (Weiss, 2001)

^d^ (Chen et al., 2010)
